# CRISPR screening by AAV episome-sequencing (CrAAVe-seq) is a highly scalable cell type-specific *in vivo* screening platform

**DOI:** 10.1101/2023.06.13.544831

**Authors:** Biswarathan Ramani, Indigo V.L. Rose, Noam Teyssier, Andrew Pan, Spencer Danner-Bocks, Tanya Sanghal, Lin Yadanar, Ruilin Tian, Keran Ma, Jorge J. Palop, Martin Kampmann

**Affiliations:** Institute for Neurodegenerative Diseases; Weill Institute for Neurosciences, University of California, San Francisco, San Francisco, CA, USA; Department of Pathology, University of California, San Francisco, San Francisco, CA, USA; Neuroscience Graduate Program, University of California, San Francisco, San Francisco, CA, USA; Biological and Medical Informatics Graduate Program, University of California, San Francisco, San Francisco, CA, USA; Biophysics Graduate Program, University of California, San Francisco, San Francisco, CA, USA; Gladstone Institute of Neurological Disease, San Francisco, CA, USA; Department of Neurology, University of California, San Francisco, San Francisco, CA, USA; Department of Biochemistry and Biophysics, University of California, San Francisco, San Francisco, CA, USA

## Abstract

There is a significant need for scalable CRISPR-based genetic screening methods that can be applied directly in mammalian tissues *in vivo* while enabling cell type-specific analysis. To address this, we developed an adeno-associated virus (AAV)-based CRISPR screening platform, CrAAVe-seq, that incorporates a Cre-sensitive sgRNA construct for pooled screening within targeted cell populations in mouse tissues. We demonstrate the utility of this approach by screening two distinct large sgRNA libraries, together targeting over 5,000 genes, in mouse brains to create a robust profile of neuron-essential genes. We validate two genes as strongly neuron-essential in both primary mouse neurons and *in vivo*, confirming the predictive power of our platform. By comparing results from individual mice and across different cell populations, we highlight the reproducibility and scalability of the platform and show that it is highly sensitive even for screening smaller neuronal subpopulations. We systematically characterize the impact of sgRNA library size, mouse cohort size, the size of the targeted cell population, viral titer, and multiplicity of infection on screen performance to establish general guidelines for large-scale *in vivo* screens.

## Main

CRISPR-based genetic screens are powerful tools for biological discovery since they enable the massively parallel interrogation of gene function. Most CRISPR screens are carried out in cultured cells. First applications in iPSC-derived brain cell types such as neurons^1,2^, microglia^3^ and astrocytes^4^ have uncovered cell type-specific mechanisms relevant to neuroscience and neurological diseases. However, a major limitation of CRISPR-based screens in cultured cells is that they do not fully recapitulate the physiological context of a multicellular organism, or tissue states such as aging, inflammation, or disease. These limitations are particularly evident in applications to biological questions related to complex organs like the brain, which involves intricate spatial interactions between numerous distinct cell types and subtypes. Therefore, *in vivo* pooled CRISPR screens directly targeting endogenous cells in the brains of mice have the potential to uncover insights that would be elusive in cell culture.

A small number of *in vivo* CRISPR screens targeting endogenous brain cells have previously been reported^5–10^, reviewed by Braun et al^11^ and summarized in **Extended Data Fig. 1a**) Some of these screens involved delivering single-guide RNA (sgRNA) libraries to the brain via lentivirus, but there is increasing recognition that lentivirus-based approaches have several drawbacks (summarized in **Extended Data Fig. 1b**). Such drawbacks include poor distribution of lentivirus through the brain and the inability to differentiate between cell types in which specific sgRNAs were expressed. To overcome some of these limitations, emerging studies have turned to adeno-associated virus (AAV), which has more widespread brain transduction, and combined with CRISPR perturbations and single-cell RNA sequencing (AAV-Perturb-Seq)^9,10^. This powerful approach provides crucial and granular details on transcriptional changes in specific cell types. However, the current cost of scaling this strategy to study larger cellular populations in the mouse brain and across multiple independent mice is prohibitive. Prior work using this approach has so far not exceeded a library size of 65 sgRNAs (targeting 29 genes) and a sampling of 60,000 cells^10^ (**Extended Data Fig. 1a**). Screens at such a scale enable the phenotypic profiling of a small number of preselected genes of interest, but not the unbiased discovery of unexpected genes that have a phenotype of interest. As such, there is a great need for a substantially more scalable *in vivo* CRISPR screening platform that retains the ability to discriminate between cell types.

We developed a strategy for screening in the mouse brain called "CRISPR screening by AAV episome sequencing" (CrAAVe-seq). Incorporating a Cre recombinase-based genetic element into the sgRNA library backbone enables the selective evaluation of phenotypes caused by genetic perturbations only in cell types of interest. Furthermore, CrAAVe-seq exploits the amplification of sgRNA sequences from AAV episomes^12,13^, rather than genomic DNA, to dramatically increase the scalability and reduce the cost of quantifying sgRNA frequencies from whole brain homogenates. Using CrAAVe-seq, we profiled neuron-essential genes in the mouse brain in different neuronal subpopulations, utilizing libraries containing ∼12,000 and ∼18,000 sgRNAs per brain and sampling at least 2.5 million neurons per brain. This approach yielded highly reproducible top hits across independent mice. Therefore, CrAAVe-seq enables high-throughput, cost-effective screening of the mammalian CNS, with immediate applicability to other cell types and tissues.

## Results

### CrAAVe-seq strategy for cell type-specific *in vivo* screening of neurons in the mouse brain

We aimed to leverage the high CNS tropism and infectivity for neurons of AAV, particularly in comparison to lentivirus, to develop an AAV-based system for pooled CRISPR perturbations of endogenous neuronal genes in the mouse brain. However, because of the ability of AAV to transduce many different cell types^14^, we sought a cell-type specific approach for CRISPR screening. To address this, we designed pAP215, an AAV vector for single-guide RNA (sgRNA) expression that contains an mU6-driven sgRNA sequence followed by a Lox71/Lox66-flanked 175 bp “handle” cassette that undergoes predominantly unidirectional^15^ inversion in cells expressing Cre recombinase (**Fig. 1a**). The construct expresses a nuclear-localized blue fluorescent protein (NLS-mTagBFP2) for visualization. A fully annotated map of the pAP215 is provided (**Supplementary File 1**).

**Fig. 1:**
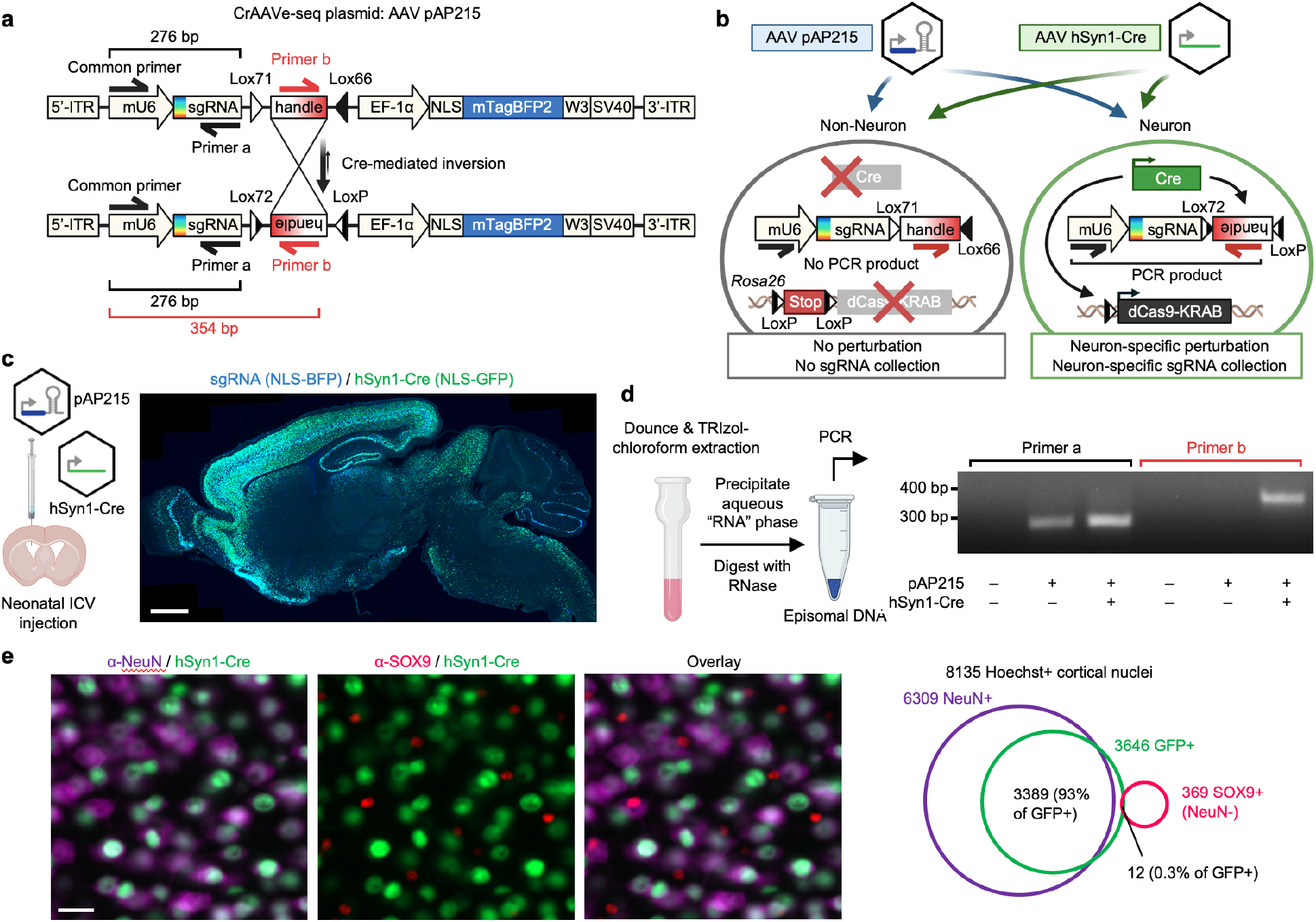
CrAAVe-seq strategy for cell type-specific *in vivo* CRISPR screening using Cre- sensitive sgRNAs. **a**, Structure of CrAAVe-seq plasmid pAP215 (sgRNA backbone), which enables cell type- specific CRISPR screening by AAV episome sequencing (CrAAVe-seq) *in vivo*. pAP215 expresses sgRNAs under an mU6 promoter and contains a Lox-flanked handle that inverts upon exposure to Cre recombinase. The plasmid also expresses a nuclear-localized BFP for visualization. **b**, Example of a strategy for CRISPRi screening in neurons using pAP215 and a Cre recombinase driven by a pan-neuronal hSyn1 promoter. Expression of Cre induces both expression of CRISPRi machinery (via recombination of LoxP sites of the Lox-Stop-Lox (LSL)- CRISPRi transgene in the endogenous *Rosa26* locus) and inversion of the pAP215 “handle” element, which contains a primer site for PCR amplification on recovered episomes. **c**, Distribution and expression of AAV pAP215 (sgRNA, nuclear BFP^+^) and AAV hSyn1-Cre-NLS- GFP (hSyn1-Cre, nuclear GFP^+^), both packaged using PHP.eB capsid, in a mouse brain three weeks after neonatal intracerebroventricular (ICV) delivery. Scale bar, 1 mm. **d**, PCR performed on episomes recovered from mouse brains expressing pAP215 with or without hSyn1-Cre using the primers diagrammed in (**a**). PCR amplification with *primer a* is invariant to Cre expression, whereas amplification with *primer b* requires Cre expression. **e**, Immunofluorescent staining for neurons (NeuN^+^) and astrocytes (SOX9^+^) in frontal cortex of mice injected with PHP.eB::hSyn1- Cre. Scale bar, 25 µm. Overlap of Cre expression with NeuN and Sox9 within a cortical region demonstrates high coverage and specificity for neurons.

We devised a strategy using pAP215 to screen for essential neuronal genes in the mouse brain *in vivo*, schematized in **Fig. 1b**. Along with uncovering key pathways involved in neuronal homeostasis, neuronal survival screens can be readily applied to various mouse models of neurological diseases, including neurodegenerative diseases, traumatic brain injury, and stroke, to identify genetic drivers or modifiers of neuronal susceptibility. AAVs containing pAP215 and promoter-driven Cre recombinase (e.g. a pan-neuronal hSyn1 promoter-driven Cre) are co-injected into the brains of inducible CRISPR interference (Lox-Stop-Lox-dCas9-KRAB, or LSL- CRISPRi) mice^16^. This leads to the activation of CRISPRi machinery and inversion of the handle sequence in neurons. PCR amplification by priming against the inverted handle on AAV episomes, followed by sequencing, enables the identification and quantification of sgRNAs in Cre-expressing neurons.

We packaged our AAV plasmids into the PHP.eB capsid, which is known to enable widespread transduction of brain cells^17^. We co-injected pAP215 (PHP.eB::pAP215) and an hSyn1 promoter-driven Cre recombinase tagged with a nuclear GFP (PHP.eB::hSyn1-Cre, diagram of construct shown in **Extended Data Fig. 1c**) by intracerebroventricular (ICV) injection into neonatal mice. We saw broad distribution of both pAP215 and hSyn1-Cre reporters across the brain, particularly in the cortex and hippocampus (**Fig. 1c**), consistent with the known distribution profile of the PHP.eB capsid^18^. Contrasting this, we observe much more limited expression and spread of lentivirus delivered by the same method and expressing the same nuclear mTagBFP reporter (**Extended Data Fig. 2**).

CRISPR screens performed in cell culture using lentivirus, which integrate their DNA into the host genome, require extraction of genomic DNA before PCR amplification of the integrated sgRNAs. In contrast, recombinant AAV genomes are mostly maintained as circular or concatenated DNA episomes, with a minor fraction integrating into the host genome^19–21^.

Consequently, a major potential advantage of AAV is that the viral genomes can be precipitated and concentrated from the aqueous phase of a TRIzol-chloroform extraction as previously demonstrated in the context of AAV capsid screening^12,13^, which could vastly improve scalability for a pooled CRISPR screen. For example, if PCR of sgRNAs from genomic DNA was necessary, a whole mouse brain weighing approximately 500 mg is expected to yield up to 1500 µg of genomic DNA. Since PCR reactions have an upper limit for the amount of template DNA (typically 10 µg per 100 µl reaction)^22,23^, screening across a single brain would require up to 15,000 µl of PCR reaction volume. This imposes a severe restriction on scalability, as large volumes of PCR reactions become economically and practically infeasible, especially for screening libraries necessitating dozens of mice.

In comparison, we performed isopropanol precipitation of nucleic acids from the aqueous phase of a TRIzol-chloroform extraction of an AAV-injected whole mouse brain. We resuspended the nucleic acid pellet in 50 µL of water and treated it with RNase, which removes all RNA without digesting the DNA of the AAV episomes. Using 1 µL of this “episome” fraction as a template, we confirmed by PCR that pAP215 could be detected in the injected brains (**Fig. 1d**). Importantly, we found that the Cre-inverted handle sequence was detectable only in mice co-injected with hSyn1- Cre (**Fig. 1d**). With the episome fraction containing a nucleic acid concentration of less than 10 ng/µL, a single 50 to 100 µL PCR reaction volume could be used if desired.

We further confirmed that PHP.eB::hSyn1-Cre expresses in neurons by immunostaining for a neuronal marker (NeuN) and an astrocyte marker (SOX9) (**Fig. 1e**). Image analysis showed that 93% of GFP^+^ co-localized with NeuN^+^ nuclei, while only 0.3% overlapped with SOX9^+^ nuclei. Of note, manual inspection of the SOX9^+^ nuclei flagged as GFP^+^ showed GFP signal was emanating from adjacent neurons that overlapped with SOX9^+^ nuclei. In effect, we did not observe any SOX9^+^ nuclei with distinct GFP signal within their nucleus, confirming hSyn1-Cre’s known selective expression in neurons^24^.

### CRISPRi knockdown *in vivo* using AAV

To test that the pAP215 plasmid is effective for CRISPRi knockdown, we used an sgRNA targeting Creb1 (sgCreb1), which encodes a ubiquitously expressed nuclear protein that is not essential for neuronal survival^25^. We also generated a non-targeting sgRNA control (sgNTC) in pAP215. We co-injected PHP.eB::pAP215-sgCreb1 or PHP.eB::pAP215-sgNTC alongside PHP.eB::hSyn1-Cre by ICV into neonatal LSL-CRISPRi mice, at approximately 1×10^11^ viral particles of each virus per mouse. We also included a group of mice injected with sgCreb1 alone to check for leakiness of the CRISPRi machinery. Three weeks after neonatal ICV injection, immunofluorescence staining confirmed strong knockdown of endogenous CREB1 in all neurons that received both sgCreb1 (BFP^+^ nuclei) and hSyn1-Cre (GFP^+^ nuclei) (**Fig. 2a-c, Extended Data Fig. 3,4**). In contrast, neurons that received sgCreb1 alone did not show knockdown. We also observed that the vast majority (>90%) of the cortex and hippocampus express both the Cre and the sgRNA, with less co-infection in striatum and cerebellum (**Fig. 2d**). This indicated that co-delivery of the sgRNA and Cre viruses can provide broad coverage and CRISPRi activity for a broad distribution of brain regions.We noted that one of the sgCreb1+hSyn1-Cre mice showed overall less viral transduction throughout the brain, most likely due to a technical issue during the injection of the virus (mouse #1). However, even with sparse infection, there was still strong knockdown of CREB1 in all BFP^+^/GFP^+^ nuclei at the level of individual cells, suggesting that transduction by an AAV at low MOI (potentially a single sgRNA) is sufficient for knockdown. Further supporting this conclusion, we found no correlation between BFP levels and the degree of CREB1 knockdown in a brain transduced with greater amounts of sgCreb1+hSyn1-Cre (mouse #3) (**Extended Data Fig. 4c**).

**Fig. 2:**
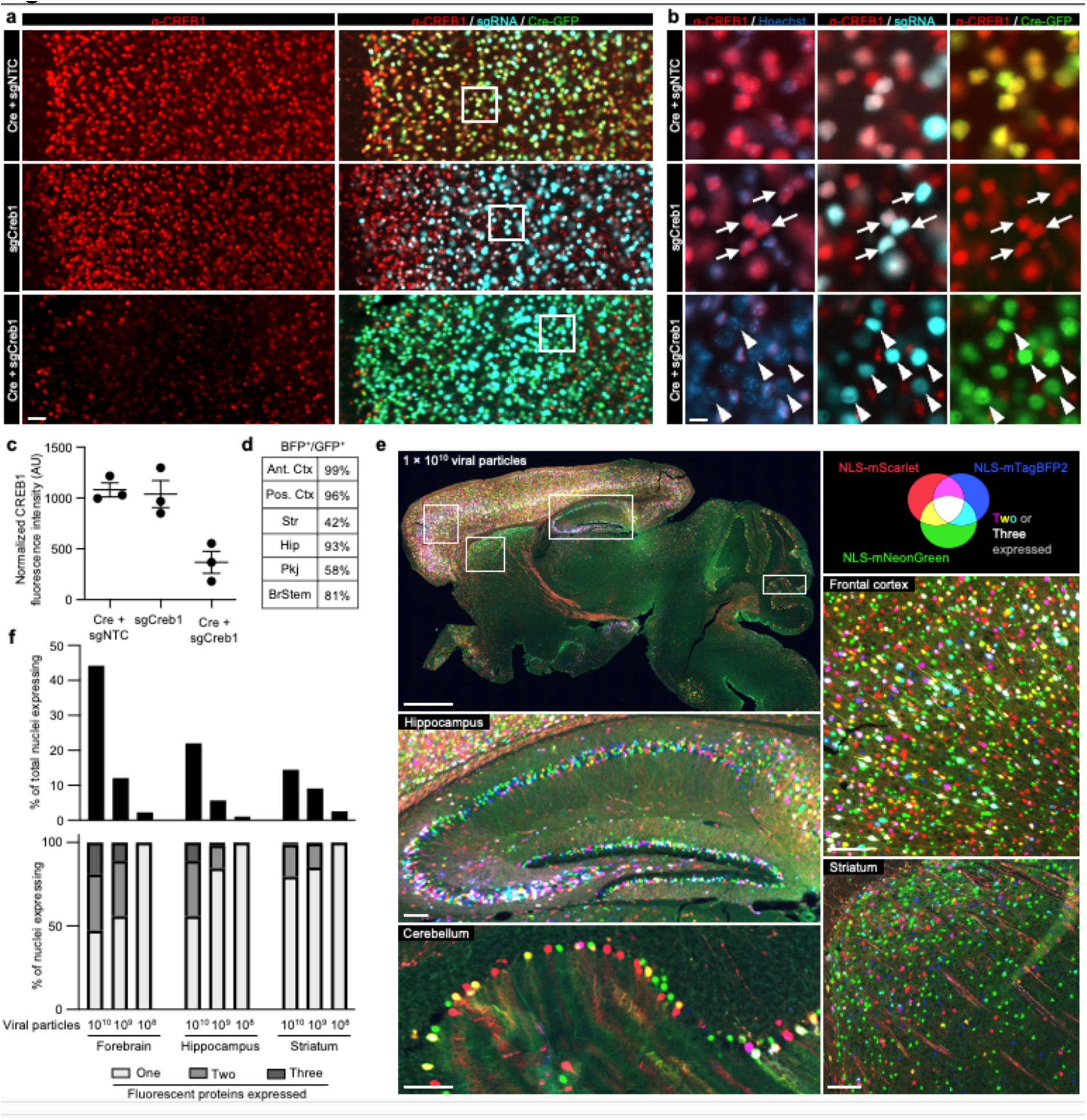
Cre-dependent CRISPRi knockdown of CREB1 *in vivo* and estimating extent of AAV multiple infections. **a**, sgRNAs targeting the non-essential gene *Creb1* (sgCreb1) or a non-targeting control (sgNTC) were cloned into pAP215 and PHP.eB-packaged, then delivered with or without PHP.eB::hSyn1-Cre by ICV injection into neonatal LSL-CRISPRi mice. Brains were stained at 3 weeks to determine reduction in CREB1 levels; a representative area of frontal cortex is shown, with cortical layer 1 oriented to the left. Scale bar, 50 µm. Other brain regions are shown in Extended Data Fig. 3. Images with higher brightness are used to identify cells with low GFP or BFP expression. **b**, Higher magnification of the boxed regions in panels from **a**, with arrowheads indicating examples of neurons having received both sgCreb1 and Cre, whereas arrows indicate neurons that received Cre only or sgCreb1 only. Scale bar, 10 µm. **c**, Quantification of CREB1 levels in sgRNA-containing (BFP^+^) nuclei within a representative region of cortex (mean ± s.e.m, n = 3 independent mice). Other brain regions are quantified in Extended Data Fig. 4. **d,** Table showing the percentage of neurons in a sgCreb1 + Cre mouse 2 that are positive for both BFP and GFP, and in different brain regions (Ctx: cortex; Pkj: Purkinje; Hippo: hippocampus; BrStem: brainstem; Str: striatum. **e**, To estimate the number of multiple infections, AAV (PHP.eB serotype) expressing nuclear-localized mNeonGreen, mTagBFP2, or mScarlet fluorescent proteins were co-packaged and co-injected by ICV into neonates at three different concentrations, and the highest concentration (1×10^10^ viral particles per mouse) is shown here across multiple brain regions. Lower concentrations are shown in Extended Data Fig. 5. **f**, *Top*, quantification of data from (**e**) and Extended Data Fig. 5, showing percentage of nuclei that were expressing at least one fluorescent protein in the forebrain, hippocampus, and striatum at three different concentrations of injected AAV (1×10^10^, 10^9^, and 10^8^ viral particles per mouse). *Bottom*, percent of nuclei expressing one, two, or all three fluorescent proteins for the same experiments.

### Distribution and extent of multiple infections by AAV

Given the high degree of co-infection between the sgRNA and Cre above in most mice, we next aimed to obtain a semi-quantitative estimate of multiplicity of infection (MOI) by injecting AAVs at multiple concentrations. We co-packaged equimolar concentrations of AAV plasmids encoding three different nuclear-localized fluorescent proteins and performed neonatal ICV injections using three different total viral particle amounts per mouse: 1×10^10^, 1×10^9^, and 1×10^8^. Three weeks post-injection, we imaged representative sagittal brain sections to evaluate the prevalence of single-, double-, and triple-infected nuclei across different brain regions.

Even at the highest concentration of AAV injected, and in the most infection-prone regions, the majority of nuclei expressed only one or two fluorescent proteins, with approximately 15% of nuclei in the densely transduced forebrain showing infection with all three fluorescent proteins (**Fig. 2e,f**). Regions with higher co-infection rates correlated directly with areas of enhanced viral tropism, such as the middle to deeper layers of the cortex and the CA3 and CA2 sectors of the hippocampus. At lower concentrations, there was a sharp decline in the total number of transduced nuclei, accompanied by a higher fraction of nuclei expressing only one type of fluorescent protein (**Fig. 2f, Extended Data Fig. 5**). In short, the results indicated that higher viral concentrations maximized the number of transduced neurons, and that even at 1×10^10^ viral particles per brain, a substantial number of cells in most brain regions express a single type of fluorescent protein, indicating that most of these cells are transduced by a single virus (with presumably a small fraction of these representing co-infection by two viruses expressing the same fluorescent protein). Obtaining a precise MOI from these experiments is challenging given the degree of variability of infection between different brain regions and neuronal types as dictated by viral tropism. Despite this limitation, these findings help us broadly estimate that the MOI across the whole brain injected with 1x10^10^ viral particles most likely ranges from 1 to 3, and could be higher than 3 in focal areas of strong viral tropism.

### CrAAVe-seq identifies neuron-essential genes

We generated two different sgRNA pooled libraries in the pAP215 AAV backbone by transferring sgRNAs from our previously established mouse sgRNA libraries^26^. This includes the 12,350- element sgRNA “M1” library that contains sgRNAs targeting 2,269 genes, including kinases, phosphatases, and other druggable targets, and the 14,975-element sgRNA “M3” library containing sgRNAs targeting 2,800 proteostasis and stress genes^26^. The M1 and M3 libraries contain 250 and 290 non-targeting sgRNAs, respectively. We co-delivered PHP.eB::hSyn1-Cre alongside either PHP.eB::pAP215-M1 library or PHP.eB::pAP215-M3 library by ICV injection into LSL-CRISPRi mice. We injected n=13 mice with the M1 library and n=19 mice with the M3 library. We collected the brains after 6 weeks, recovered episomes and amplified sgRNAs using primers specific to the Cre-inverted handle in pAP215, followed by next-generation sequencing (**Fig. 3a**). We compared the sgRNA frequencies from the recovered episomes to the sgRNA frequencies of the packaged and injected AAV sgRNA libraries to identify changes in sgRNA frequency. We refer to this workflow as “CrAAVe-seq" throughout the rest of this manuscript. The age, sex, and sgRNA read-depth for each mouse is provided in **Supplementary Table 1**.

**Fig. 3:**
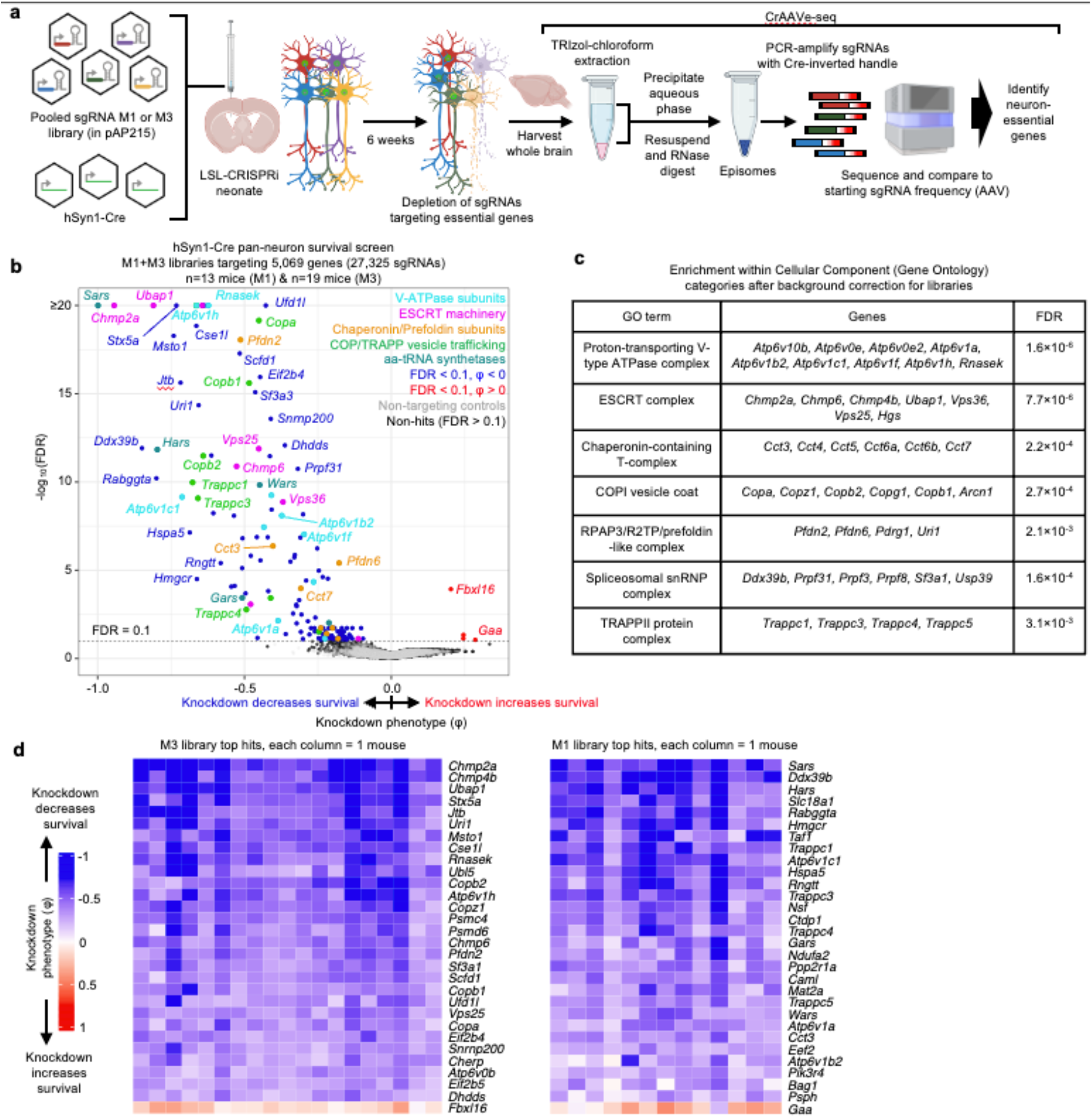
*In vivo* CrAAVe-seq uncovers neuron-essential genes in the mouse brain. **a**, Experimental design for CrAAVe-seq to identify neuron-essential genes across two different pooled sgRNA libraries (pAP215 expressing the M1 or M3 libraries). M1 targets kinases, phosphatases, and druggable targets: 2,269 genes (12,100 targeting sgRNAs and 250 non- targeting controls); M3 targets stress and proteostasis targets: 2,800 genes (14,685 targeting sgRNAs and 290 non-targeting controls)^26^. The sgRNA libraries and hSyn1-Cre were packaged into AAV (PHP.eB capsid) and co-injected into neonatal LSL-CRISPRi mice. **b**, Results from CRISPRi screen described in (**a**): Knockdown phenotype (φ, see Methods) for 5,069 genes and non-targeting (quasi-gene) controls at 6 weeks post-injection (n=13 mice injected with pAP215- M1 library, and n=19 mice injected with pAP215-M3 library). Hit genes (FDR<0.1) within enriched pathways are color-coded. Non-targeting controls (gray) are plotted on top of non-hit genes (black). **c**, Table showing enrichment of hit genes in different Cellular Component Gene Ontology (GO) terms. **d**, Knockdown phenotype for each individual mouse (columns) for 30 top hit genes (rows, 29 with negative and 1 with positive knockdown phenotypes). Rows are ranked by average knockdown phenotype.

Analysis of changes in sgRNA frequencies using an optimized computational pipeline (see Methods for details) identified 147 genes with significant sgRNA depletion (false discovery rate (FDR) < 0.1), indicating that knockdown of these genes promoted neuronal death (**Fig. 3b**). The output knockdown phenotype of all screens is provided in **Supplementary Table 2**. Gene-set enrichment analysis (with background correction for the genes in the library) further highlighted that many hit genes belonged to distinct biological categories (**Fig. 3c**). Top categories included members of the vacuolar ATPase complex, ESCRT pathways, COP-I and COP-II vesicle trafficking, and members of the chaperonin/TRiC complex. Although not identified from gene enrichment, we also noted some of the top hits to be members of the aminoacyl tRNA synthetase families. Supporting the notion that the genes with a negative knockdown phenotype are neuron-essential genes, we found that 91 of the 147 hit genes (62%) foverlapped with known common essential genes established by DepMap from cancer cell lines^27^ (**Extended Data Fig. 6a**). We similarly found substantial overlap with our prior screens for essential genes performed in induced pluripotent stem cell-derived neurons^2^ (**Extended Data. Fig. 6b**). Even though the vast majority of our hits are known essential genes in human cancer cell lines or human iPSC-derived neurons, this method reveals some unique hits that have not been previously reported, including *Jtb, Snx17, Psph, Bag1*, and *Becn1*. We also note the presence a few genes for which the sgRNAs were weakly, but significantly enriched relative to input AAV, suggesting that knockdown of these genes enhances neuronal survival.

To examine the robustness of our screening approach, we compared knockdown phenotypes for top hits from each library in each mouse and found that phenotypes were highly reproducible across individual mice (**Fig. 3d**), corroborating that hits obtained from analyzing the full cohort of mice are not entirely driven by a subset of mice. We also compared two individual mice each from the M1 and M3 sgRNA library cohorts, selecting the mice that called the highest number of hits. We again observed a strong correlation in knockdown phenotypes, particularly for the strongest hit genes (**Extended Data Fig. 7a**). The knockdown phenotypes also strongly correlated when comparing male versus female mice from each cohort; no sex-specific neuron-essential genes were observed (**Extended Data Fig. 7b**). We further confirmed that the knockdown phenotypes of the top hit genes in the M1 library require the expression of hSyn1-Cre (**Extended Data Fig. 8a)**. The knockdown phenotypes for all genes per individual mouse are provided in **Supplemental Table 3**.

The amount of injected AAV library for the M1 and M3 screens above were 2×10^10^ and 5×10^10^ viral particles, respectively, per mouse. Based on our MOI experiments (**Fig. 2d,e**), these concentrations are expected to transduce neurons in several brain regions with an MOI of greater than 1, leading to the expression of multiple different sgRNAs within some neurons. To investigate the impact of MOI on screen quality, we injected a cohort of LSL-CRISPRi littermates with two lower concentrations of the M1 AAV sgRNA library alongside a constant amount of hSyn1-Cre (1×10^11^ viral particles per mouse). Evaluating n=2 mice injected with an intermediate amount (7×10^9^ viral particles) of the library showed a clear, strong correlation of the sgRNAs frequencies recovered from mouse brains with those in the input AAV library (**Extended Data Fig. 8b).** In contrast, n=4 mice injected with a 10-fold lower concentration (7×10^8^ viral particles) showed much poorer correlation and extensive sgRNA dropout. The intermediate concentration identified 20 hits that overlap with hits of the M1 sgRNA library cohort from Fig. 3b, while the lower concentration identified just 2 hits meeting these criteria (**Extended Data Fig. 8c**). In summary, although the concentrations used for the original screens have an MOI greater than 1 in several brain regions, injecting smaller amounts of virus dramatically reduces the power of the screen. We suggest that the poor performance of lower viral concentrations is due to reducing coverage, indicated by vastly increased sgRNA dropout in the lower concentration. Conversely, the infection of a proportion of cells with more than one sgRNA expected at higher concentrations seems to have a much less detrimental impact on the screen quality, likely because the majority of sgRNAs does not cause a phenotype, and because each individual sgRNA is screened in many independent neurons.

### Screening for essential genes in CaMKII^+^ neurons *in vivo*

To determine if a different neuronal Cre driver targeting a large neuronal subpopulation produces similar findings, we tested a CaMKII promoter that has traditionally been used to target predominantly forebrain excitatory neurons^28^. We first co-injected PHP.eB::CaMKII-Cre, a LoxP-dependent GFP reporter (FLEX-GFP), and PHP.eB::pAP215-sgCreb1 into an LSL-CRISPRi mice. We observed widespread FLEX-GFP expression throughout the brain, particularly strong in the cortex (**Fig. 4a**). This broad promoter activity is consistent with recent reports of AAV-delivered CaMKII-driven reporters in the mouse brain^29,30^. In areas of the forebrain with high GFP expression, we observed strong knockdown of CREB1 in BFP^+^ nuclei (**Fig. 4a,b**). We performed CrAAVe-seq in cohort of 12 mice injected with PHP.eB::pAP215-M1 library and CaMKII-Cre, revealing a robust profile of neuron-essential genes (**Fig. 4c**). The knockdown phenotypes of the top hits of this CaMKII-Cre cohort correlated very strongly with the hits from the hSyn1-Cre cohort of Fig. 3 (**Fig. 4d**). This further supported that CrAAVe-seq reproducibly detects neuron-essential genes *in vivo*.

**Fig. 4.**
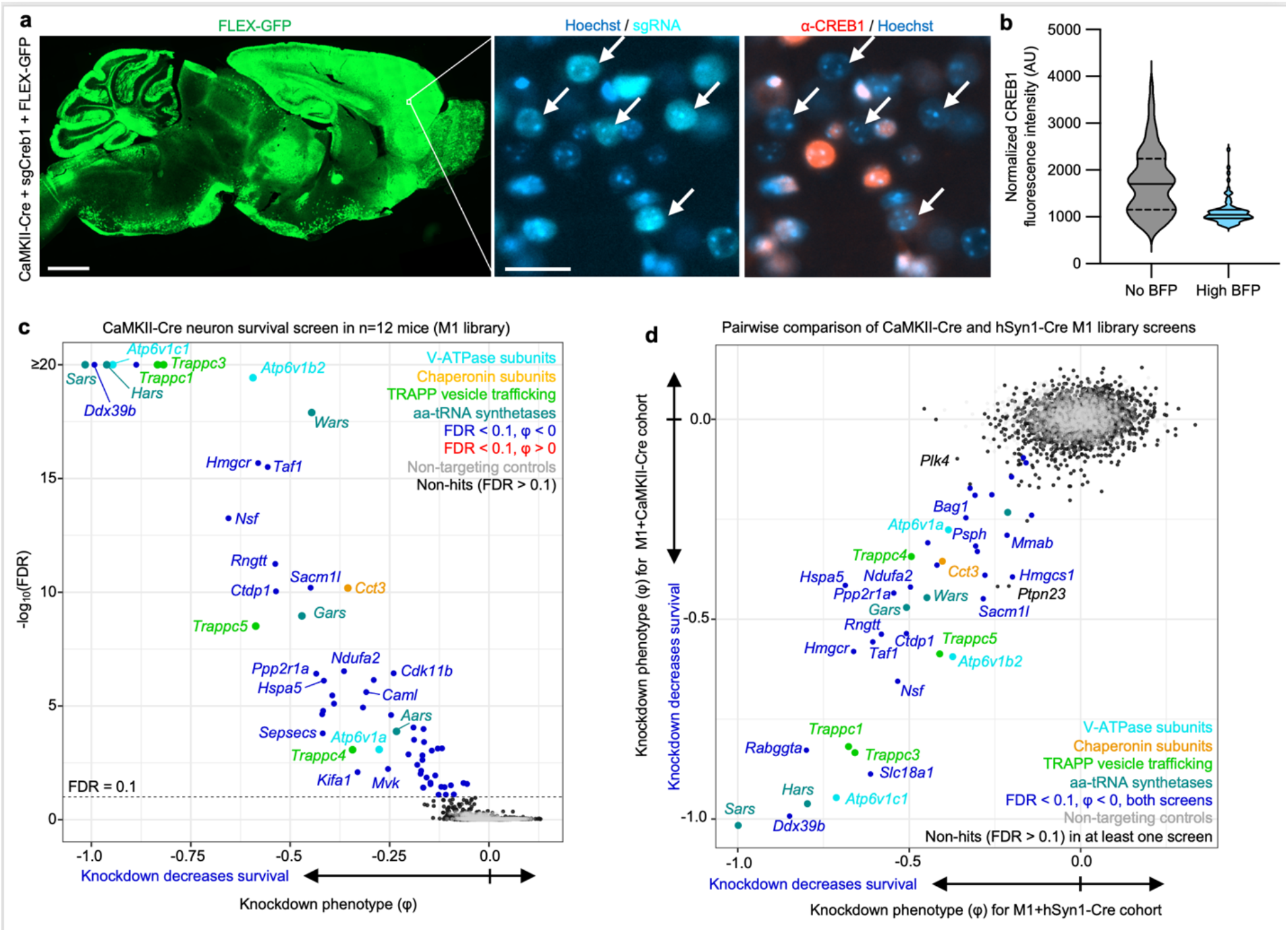
Genetic modifiers of survival in a CaMKII^+^ subpopulation of neurons in the mouse brain *in vivo*. **a**, PHP.eB::CaMKII-Cre was co-injected with PHP.eB::FLEX-GFP and PHP.eB::pAP215- sgCreb1 in LSL-CRISPRi mice. Brains were examined by immunofluorescence staining for CREB1 at 3 weeks. *Left*, low-power view showing expression of FLEX-GFP across a sagittal section of the brain. *Right*, magnified region of frontal cortex showing CREB1 knockdown in BFP^+^ (sgCreb1^+^) cells (arrows). **b,** CREB1 levels in BFP^+^ (n=79 cells) versus BFP^-^ (n=78 cells) nuclei in a representative region of cortex. Scale bars: 1 mm (left) and 20 µm (right). **c**, Knockdown phenotypes for 2,269 genes averaged across 12 mice at 6 weeks after ICV injection of PHP.eB-packaged M1 sgRNA library and CaMKII-Cre. Experimental design is identical to Fig. 3a, except injecting CaMKII-Cre instead of hSyn1-Cre. Genes within enriched pathways are color-coded. **d,** Comparison of CRISPRi screen results between the CaMKII-Cre cohort (**c**) and the hSyn1-Cre cohort shown in Fig. 3b.

### Screening on small neuronal subpopulations and requirement of the Cre-inverted handle

We next wanted to evaluate whether CrAAVe-seq is sensitive enough to identify neuron-essential genes from smaller neuronal subpopulations. We selected a recently developed AAV plasmid that utilizes enhancer elements designed to express Cre recombinase in forebrain GABAergic neurons (CN1851-rAAV-hI56i-minBglobin-iCre-4X2C-WPRE3-BGHpA, which we refer to here as “hI56i-Cre”)^31^. Co-injection of this Cre alongside PHP.eB::pAP215-sgCreb1 and PHP.eB::FLEX-GFP in LSL-CRISPRi mice showed its distribution to be much more restricted than that of hSyn1- Cre and CaMKII-Cre, with a large degree of transduction within the olfactory bulb (**Fig. 5a**). In areas of the cortex with sparse GFP expression where individual non-overlapping nuclei could be confidently examined, GFP^+^ nuclei showed reduced levels of CREB1, again confirming Cre- dependent CRISPRi knockdown (**Fig. 5b**).

**Fig. 5:**
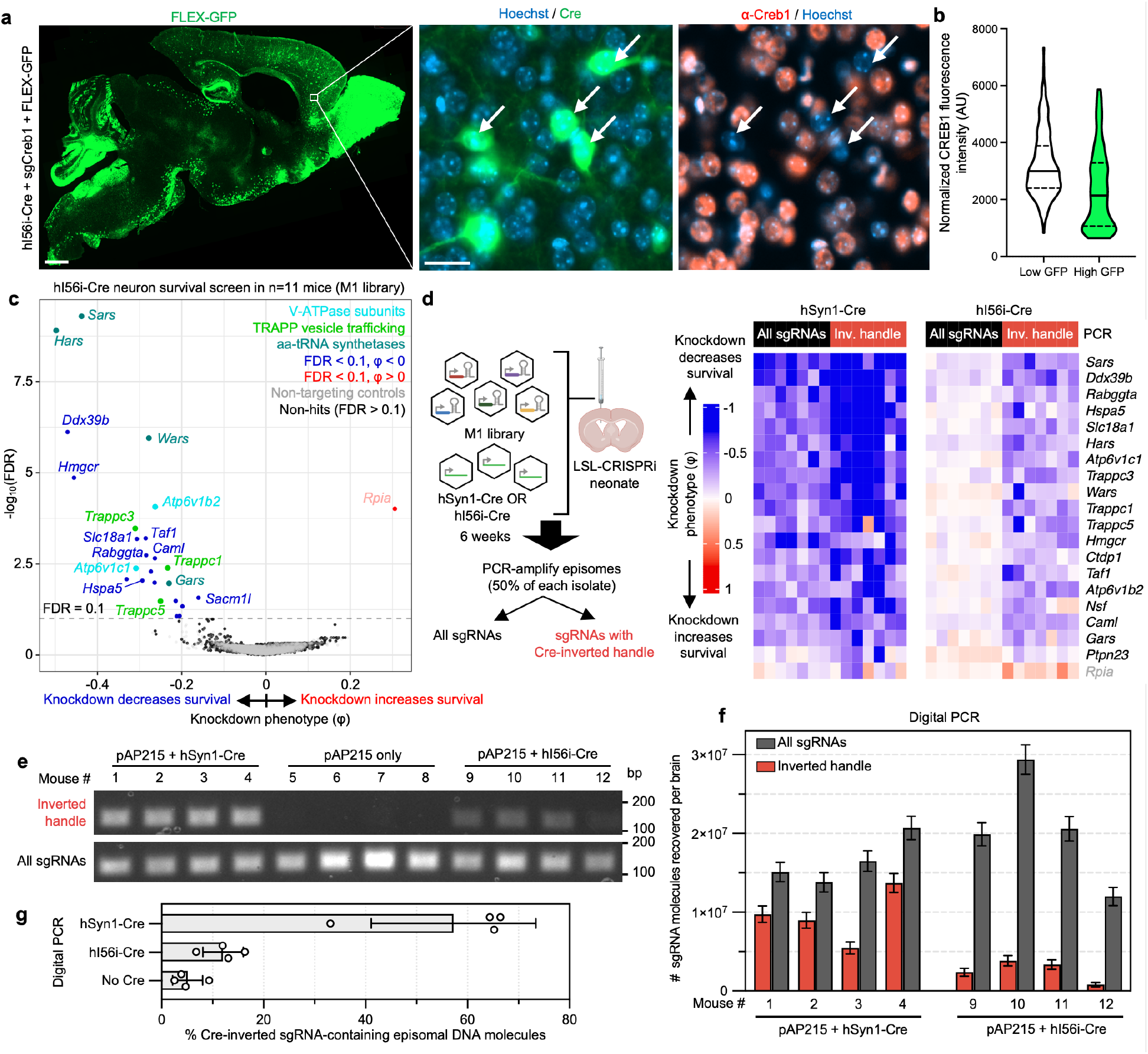
Cre-dependent sgRNA recovery is critical for screening on small neuronal subpopulations. **a**, PHP.eB::hI56i-Cre (predicted to target predominantly forebrain GABAergic interneurons)^31^ was co-injected with PHP.eB::FLEX-GFP and PHP.eB::pAP215-sgCreb1 in LSL-CRISPRi mice, followed by immunofluorescence staining for CREB1 in the brain at 3 weeks. *Left*, Sagittal brain section showing FLEX-GFP expression (reporting Cre activity). *Right*, Inset showing CREB1 knockdown in GFP^+^ cells (arrows). Scale bars: 800 µm (left) and 20 µm (right). **b**, CREB1 levels in GFP^+^ (n=189 cells) versus GFP^-^ (n=188 cells) cells within a representative cortical region**. c**, Averaged knockdown phenotypes for 2,269 genes across 11 mice, 6 weeks after ICV injection of PHP.eB::M1 sgRNA library and PHP.eB::hI56i-Cre. *Rpia* is labeled in light pink, indicating its phenotype was driven by only one sgRNA. **d**, *Left*, Experimental design comparing impact of PCR recovery of all sgRNAs versus only those with the Cre-inverted handle, in mice injected with M1 library and either hSyn1-Cre or hI56i-Cre. *Right*, Heatmap showing knockdown phenotypes for top hits (rows) across n=7 mice (columns) from each Cre cohort, contrasting the two PCR recovery methods. Genes listed are top hits selected from (**c**) and include 19 with negative and 1 with positive knockdown phenotypes. **e**, Ethidium bromide-stained agarose gel for PCR products from episomes recovered from brains of mice at 4 weeks post-ICV injection. **f,** Digital PCR (dPCR) performed on Cre-injected samples in (**e**) showing absolute numbers of pAP215 sgRNA-encoding episomal DNA molecules recovered from each brain using CrAAVe- seq (with 95% confidence interval from dPCR Poisson distribution), both for total episomes and episomes with cell type-specific Cre-inverted handle. **g**, Percent recovered sgRNAs from (**f**) that contains the inverted handle upon hSyn1-Cre (all neurons) or hI56i-Cre (small subset) expression, including no Cre negative control (n=4 mice per condition, mean ± s.d.).

**Fig. 6:**
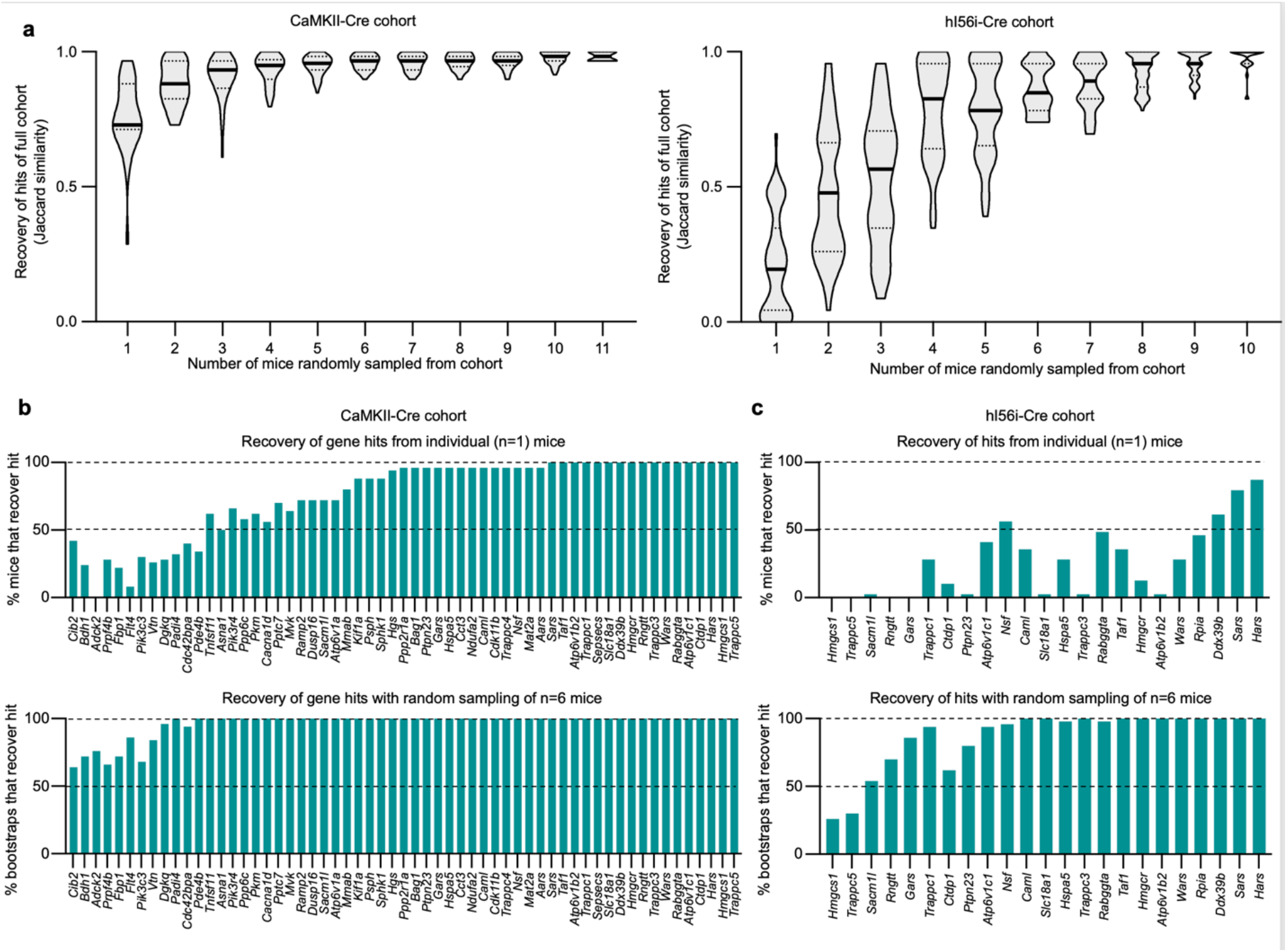
Bootstrapping analysis estimates the number of mice required for different *in vivo* screening conditions. **a**, Increasing numbers of mice were randomly sampled and bootstrapped from the CaMKII-Cre and hI56i-Cre cohorts to assess how frequently the hits from a bootstrap overlap with the hits identified from their full respective cohort. Hits are defined by an FDR < 0.1. **b**, *Top*, the percentage of individual mice that recover each of the hits defined from analyzing the full cohort. *Bottom,* percentage of bootstraps when randomly sampling 6 mice of the cohort that recover each of the hits of the full cohort.

We performed CrAAVe-seq on a cohort of 11 mice at 6 weeks after neonatal ICV injection of the PHP.eB::pAP215-M1 library alongside hI56i-Cre. This screen again revealed a profile of essential neuronal genes (**Fig. 5c**), with hits that almost fully overlapped with the hSyn1-Cre and CaMKII- Cre screens (**Extended Data Fig. 9a**). One exception was *Rpia*, which was uniquely identified as a hit in this hI56i-Cre cohort, but we also noted that its effect was driven by one sgRNA out of the 5 targeting the gene; all other hit genes resulted from at least 2 significant differentially abundant sgRNAs for that gene. Nonetheless, these screens established that CrAAVe-seq sensitively detects neuron-essential genes on a smaller neuronal population. Top hits showed consistently reproducible knockdown phenotypes in individual mice (**Extended Data Fig. 9b**).

To determine if the Cre-dependent recovery of sgRNA sequences is required for screening *in vivo*, we examined cohorts of mice injected with the PHP.eB::pAP215-M1 library together with either PHP.eB::hSyn1-Cre or PHP.eB::hI56i-Cre. 6 weeks after ICV injection, we performed episome isolation as above but used half of the episome preparation for PCR amplification of all sgRNAs, and the other half for PCR amplification of only those that underwent Cre-dependent inversion of the handle (**Fig. 5d**, primer pairs as schematized in Fig. 1a). We found that for screens conducted in large neuronal populations accessed with hSyn1-Cre expression, analyzing all sgRNAs provides similar knockdown phenotypes of the top hit genes, but also demonstrates a noticeable increase in the knockdown phenotypes for most genes when analyzing only sgRNAs with the Cre-inverted handle (**Fig. 5d**). This nevertheless indicated that hSyn1-Cre drives sgRNA activity in a sufficiently large number of neurons to permit identification of hit genes when evaluating all recovered sgRNAs, but that the presence of non-transduced or transduction in non- neuronal cells could partially mask the effects of active hits. More importantly and in contrast, screens in a small neuronal subpopulation accessed by hI56i-Cre showed that knockdown phenotypes of the top known hits required amplification with the Cre-inverted handle (**Fig. 5d**). Indeed, hit genes were significantly different compared to non-targeting controls when examining sgRNAs with the Cre-inverted handle (p < 10^−46^), but not when examining all sgRNAs (p = 0.72) (Kolmogorov-Smirnov test, **Extended Data Fig. 9c**). This supports that sgRNAs expressed in Cre-negative neurons can fully obscure the signal of active sgRNAs, making the Cre-inverting handle in pAP215 essential for screening smaller subpopulations of a broadly transduced population.

To further quantity the proportion of the total sgRNAs and Cre-inverted sgRNAs, as well as to determine their relative proportions after inversion with different Cres, we conduced PCR with different primer pairs. Episomes were recovered from mice injected PHP.eB::pAP215-M1 library alone, or with either PHP.eB::hSyn1-Cre or PHP.eB::hI56i-Cre. Standard PCR with examination of the products by gel electrophoresis showed that with primers targeting total sgRNAs, there is qualitatively an approximately equal intensity of the PCR products. When using primers targeting the Cre-inverted handle, there was a strong band in mice injected with hSyn1-Cre, no detectable product in mice without Cre, and a weak band with hI56i-Cre (**Fig. 5e**).

To determine the absolute number of template molecules between conditions, we performed digital PCR using the same primer pairs and the same samples. We first determined the total number of sgRNA molecules and found that 15–30 × 10^6^ episomal DNA molecules can be recovered from each brain, and within similar ranges in both Cre conditions (**Fig. 5f**). We then found that the number of sgRNAs containing the Cre-inverted handle in the hSyn1-Cre condition were generally a majority fraction of the total, with 5.4–13 × 10^6^ molecules detected corresponding to 57% of total sgRNAs on average (**Fig. 5f,g**). In contrast, with hI56i-Cre expression we detected far fewer molecules, 0.8–3.8 × 10^6^ molecules containing the inverted handle, corresponding to 12% of the total sgRNAs (**Fig. 5f,g**), consistent with findings on the DNA gel in **Fig. 5e**. In comparison, the no-Cre controls generated around 5% signal for inverted PCR primer pair, which we interpret as technical background, as no Cre was present within these mice to invert the handle sequence.

Overall, these results indicate that CrAAVe-seq is extremely sensitive and is critical for screening on neuronal subpopulations. Moreover, the number of molecules detected also demonstrate the substantially large numbers of neurons being sampled per mouse. Assuming at minimum, when performing CrAAVe-seq using hSyn1-Cre, we estimate from the above digital PCR experiments that 5 million Cre-inverted sgRNA molecules are captured and sequenced from a whole brain. Based on the frequency of the multiple infections observed in Fig. 2, we estimate that most of the transduced neurons express between 1 to 3 transgenes. Therefore, we estimate at least 1.67 million neurons are screened per brain. Importantly, this number can be dramatically boosted by increasing the number of mice for a screen at minimal additional cost. For example, the M3 library- injected cohort of 19 mice represents screening of at least 30 million neurons.

### Estimating number of mice required for screening in differing cell populations

To investigate the impact of cohort size on hit recovery under different Cre conditions, we performed a bootstrap analysis on the CaMKII-Cre screen cohort (from Fig. 4c) and hi56i-Cre screen cohort (from Fig. 5c), which were all injected with the same preparation and concentration of PHP.eB::pAP215-M1 library. Briefly, we randomly subsampled mice without replacement at incrementing cohort sizes 50 times for each cohort. We measured the overlaps of hits identified in the subset of mice at an FDR cutoff of 0.1 with the set of hit genes identified from the full corresponding cohort (i.e. n=12 CaMKII-Cre mice or n=11 hI56i-Cre mice) (**Fig. 6a**).

Using the cohort of mice from the CaMKII-Cre screen, we observed that the complete set of hits was recovered with fewer than the complete set of mice (n = 11); 90% of the hits were recovered with just three mice (**Fig. 6a**). When examining each of the 61 hits identified from the full CaMKII- Cre cohort, the majority of genes were called hits even in individual mice. (**Fig. 6b, *top***). When bootstrapping by randomly sampling n=6 mice from the cohort, nearly all hit genes were recovered in the vast majority of bootstraps. (**Fig. 6b, *bottom***).

In contrast, the hI56i-Cre screen cohort overall showed fewer hits, further supporting that targeting a neuronal subpopulation reduces the power of the screen (**Fig. 6a**). Moreover, while few hits were robustly recovered in individual mice (**Fig. 6c, *top***), there was a much stronger benefit of additional mice, since most hits were recovered robustly when bootstrapping on n=6 mice. (**Fig. 6c, *bottom***).

Together, these findings confirm that relatively few mice are required when screening on large neuronal populations, while more mice are required to effectively power screens when screening on neuronal subpopulations. In future applications, this bootstrapping approach can be implemented to empirically establish whether an *in vivo* screen is powered for the given neuronal population and library size.

### Screening on essential chaperone genes with a focused sgRNA library

To test the impact of using a smaller sgRNA library on screening performance by CrAAVe-seq, we designed a library of 2,172 sgRNAs targeting 354 known molecular chaperone and co-chaperone genes and cloned it into the pAP215 vector (pAP215-chap library). Given the importance of protein homeostasis in neurons, this would also further establish the chaperones that are critical for neuronal survival. We co-injected PHP.eB::pAP215-Chap library and PHP.eB::hSyn1-Cre into a cohort of 4 neonatal LSL-CRISPRi mice, followed by CrAAVe-seq at 6 weeks. The screen revealed 32 hit genes, 26 of which had a negative knockdown phenotype indicating them to be neuron-essential chaperones (**Fig. 7a**). The hits included several genes that were identified from the M1 and M3 library screens in Fig. 3 (e.g. *Hspa5* and subunits of the chaperonin complex). Notably, contrasting the M3 library screen, the chaperonin subunits were even more strongly represented and exhibited a lower FDR, supporting that a reduced library size increased the power of this screen. The high power and sensitivity of this screen with a focused library is further corroborated by the finding that more than half of the 32 hits could be captured from one mouse, and this number substantially increases with using 2 mice (**Fig. 7b**).

**Fig. 7:**
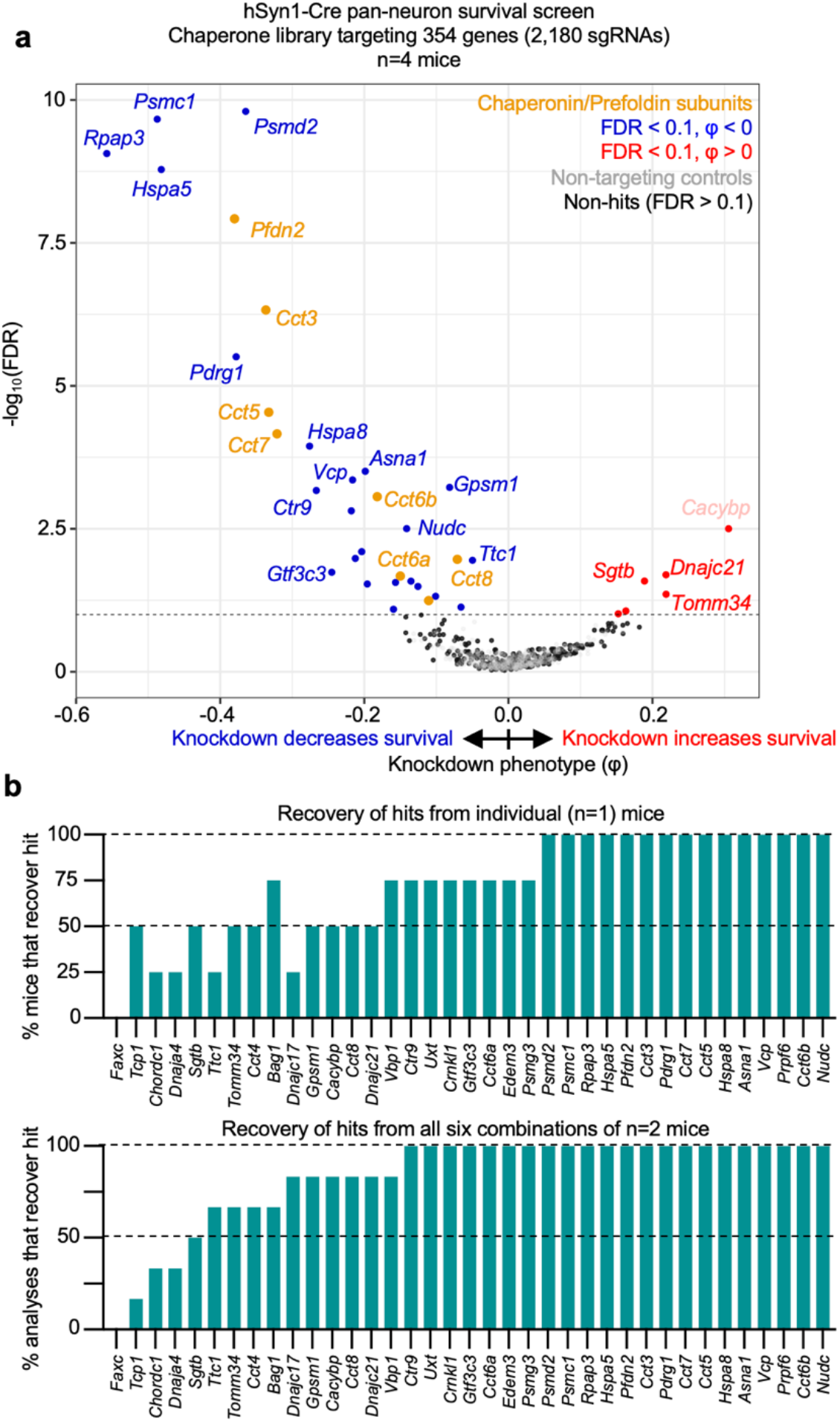
CRISPRi screening for neuron-essential chaperones using a smaller sgRNA library. **a,** Knockdown phenotypes of 354 chaperone genes averaged across n=4 LSL-CRISPRi mice at 6 weeks after neonatal ICV injection of PHP.eB::hSyn1-Cre and a PHP.eB-packaged sgRNA library targeting chaperones (354 genes, 2,180 sgRNAs, of which 350 are non-targeting controls). *Cacybp* is labeled in light pink to reflect that its phenotype was driven by only one sgRNA. **b**, *Top*, the percentage of individual mice that recover each of the hits defined from analyzing the full cohort. *Bottom,* percentage of analyses for all six combinations of n=2 of the 4 mice that recover each of the hits of the full cohort.

### Confirmation of *Hspa5* and *Rabggta* as essential genes in mouse neurons

We selected two top hits from our *in vivo* screens, *Hspa5* and *Rabggta*, for individual validation to establish that screening for sgRNA depletion from recovered episomes reflects capture of neuron-essential genes. *Hspa5* and *Rabggta* were also consistently among the strongest hits in individual mice and highlight two distinct biological pathways. *Hspa5* encodes an ER-associated Hsp70 chaperone, helping facilitate the folding of newly synthesized proteins. Prior work has shown that knockdown or knockout of *Hspa5* in select neuronal populations *in vivo* leads to their apoptotic cell death^32,33^. *Rabggta* encodes a geranylgeranyl transferase that regulates GTPase activity of Rab proteins, which are involved in intracellular vesicle trafficking. The impact of knocking down these genes broadly in neurons had not been previously examined.

In primary neurons cultured from mice with constitutive CRISPRi machinery, we confirmed that sgRNAs targeting *Hspa5* (sgHspa5) and *Rabggta* (sgRabggta) suppress expression of their respective endogenous transcripts (**Fig. 8a, f**). Furthermore, injection of PHP.eB::pAP215- sgHspa5 and PHP.eB::hSyn1-Cre into neonatal LSL-CRISPRi mice led to a severe motor phenotype after approximately 2 weeks in mice, but not the sgRNA alone (**Supplementary Videos 1 and 2**), requiring prompt euthanasia. The brains from mice co-injected with sgHspa5 and hSyn1-Cre were markedly smaller in size relative to littermates with sgHspa5 alone (**Fig. 8b,c**). In primary neurons cultured from LSL-CRISPRi mice, AAVs delivering sgHspa5 led to marked Cre-dependent neuronal death within 2 weeks of expression (**Fig. 8d,e**).

**Fig. 8:**
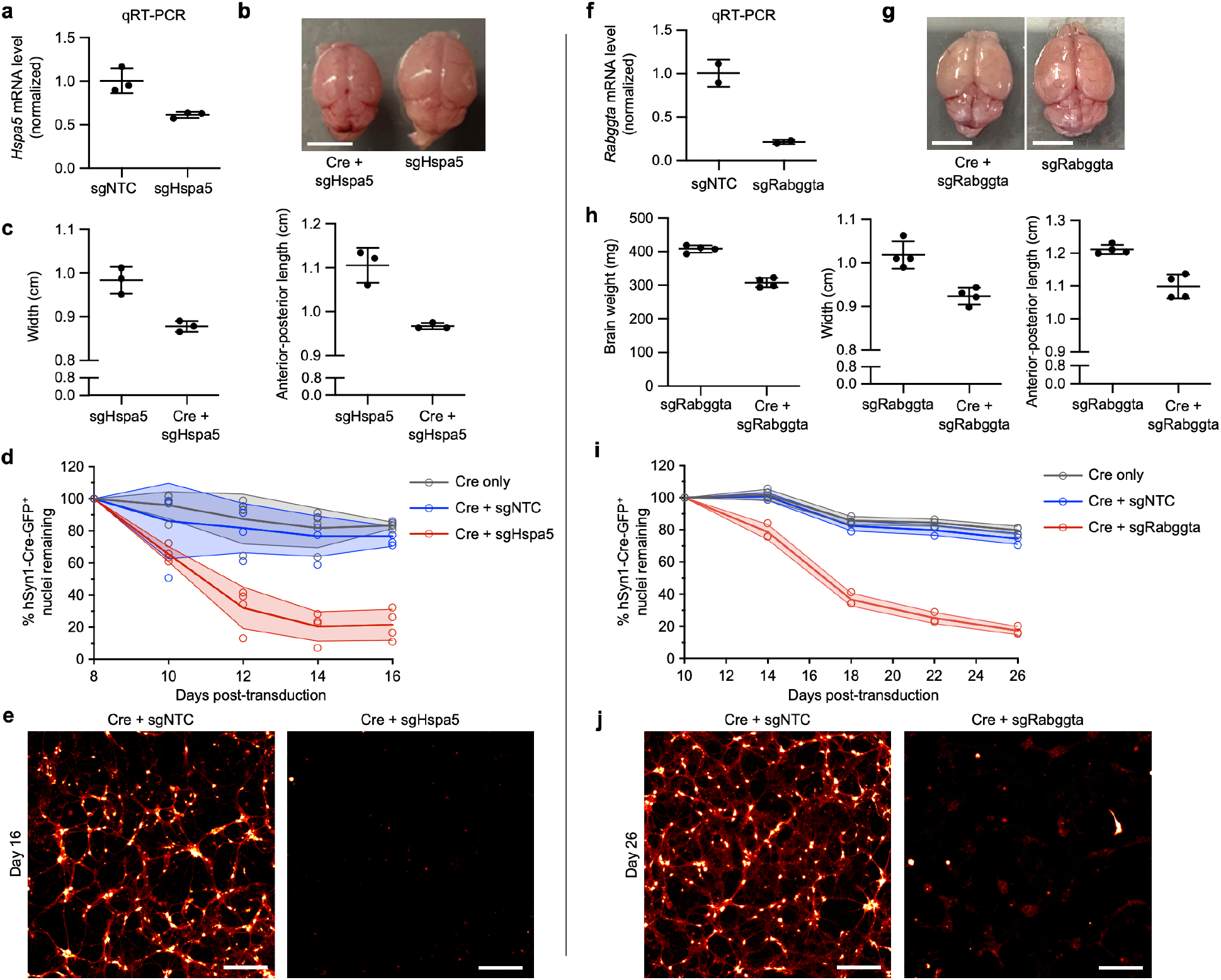
Validation of *Hspa5* and *Rabggta* as neuron-essential genes in mouse neurons. **a**, Primary neurons cultured from constitutive CRISPRi mice were transduced with PHP.eB::pAP215 targeting *Hspa5* (sgHspa5) or non-targeting control (sgNTC). *Hspa5* mRNA levels were assayed by qRT-PCR and normalized to the sgNTC control (mean ± s.d., n=3 technical replicates). **b**, Representative brains of LSL-CRISPRi mice 16 days after neonatal ICV injection of PHP.eB::pAP215-sgHspa5 with or without co-injection with PHP.eB::hSyn1-Cre (Cre). Scale bar: 5 mm. **c**, Quantification of brain width and length (mean ± s.d., n = 3 independent mice) from (**b**). Gross motor phenotypes for these mice are shown in Supplementary Videos 1 and 2. **d**, Primary neurons cultured from LSL-CRISPRi mice following transduction with PHP.eB::hSyn1-Cre and PHP.eB::pAP215-sgHspa5 or sgNTC (mean ± s.d., n = 4 wells). Survival was determined by counting GFP+ nuclei over time and normalized to peak fluorescence, which occurred at day 8. **e**, Representative image of primary neuronal cultures from (**d**) at 16 days post transduction. Cells were co-transduced with constitutively expressed cytosolic mScarlet (PHP.eB::CAG-mScarlet), which reveals fine neuronal processes (displayed with a red to white lookup table). Scale bar: 250 µm. **f-j**, Identical experimental designs to (**a-e**), except using PHP.eB::pAP215-sgRabggta instead of sgHspa5, with the following changes: Brains extracted from mice in (**h**) were weighed after extraction. Quantification of neuronal survival in (**i**) used n=3 wells instead of n=4. Peak fluorescence in cultures quantified in (**i**) occurred at day 10 instead of day 8, and were normalized to that timepoint, and imaged until day 26 instead of day 16. Neurons in (**j**) were displayed at day 26 instead of day 16.

Similarly, neonatal LSL-CRISPRi mice co-injected with sgRabggta and hSyn1-Cre showed a severe motor phenotype at 24 days after injection. Following euthanasia, the brains of these mice weighed less and measured smaller in comparison to littermates injected with sgRabggta alone (**Fig. 8g,h**). Primary neurons transduced with sgRabggta and hSyn1-Cre showed increased neuronal death beginning at approximately 14 days after transduction, with near complete death at 26 days (**Fig. 8i,j**). These findings confirm that *in vivo* CRISPR screening using CrAAVe-seq identifies essential genes that validate even when sgRNAs are introduced into matured primary neuronal cultures postnatally.

## Discussion

Here we established CrAAVe-seq, a platform for cell type-specific CRISPR-based screening *in vivo* in mouse tissues with very high scalability due to the amplification of sgRNA sequences from AAV-derived episomes. Our pan-neuronal CRISPRi screen targeting over one quarter of the protein-coding genes in the mouse genome uncovered neuron-essential genes in the brain with high reproducibility of top hits between individual mice. Many of these genes have been previously recognized as common essential genes by DepMap^27^ or in our prior screens in iPSC neurons^2^. Some hits were unique to the *in vivo* screen, and include *Jtb* (involved in cytokinesis), as well as *Snx17 and Snx20,* members of the sorting nexin family. Therefore, our screens already begin to uncover neuronal vulnerability genes in the mouse brain that have not been previously documented. Conversely, some hits were strong essential genes in our screens in mono-cultured human iPSC-derived neurons, such as *SOD1* and genes involved in the exocyst complex (*EXOC3, EXOC7*), possibly because they cause vulnerabilities that are buffered in the context of the brain. Our screens also uncovered a few genes whose knockdown produced a positive phenotype, and include *Fbxl16*, *Gaa*, *Dstn*, and *Kdelc16*. It is unclear whether these gene knockdowns increase resiliency from death or from inducing proliferation of a transduced cell population; they warrant further investigation.

Our Cre-dependent sgRNA recovery strategy enables highly sensitive screens in smaller neuronal subpopulations, providing an avenue for CRISPR screening on molecularly defined cell types that can be accessed through specific Cre-drivers, or brain regions that can be physically dissected. One area for ongoing optimization is increasing sgRNA recovery and coverage across cell populations. Our current proof-of-concept studies rely on co-injecting AAVs for the sgRNA libraries and Cre recombinase, but using transgenic Cre mouse lines could maximize active sgRNA expression without relying on co-infection. Additionally, temporal control over gene perturbation can be achieved by delivering sgRNA libraries later, via intravenous delivery, or by crossing LSL- CRISPRi mice to Cre^ERT2^ lines for tamoxifen-inducible activation. Furthermore, the Lox71/Lox66 system used for unidirectional handle inversion may not be the maximum possible efficient, so we are developing a FLEX-based AAV sgRNA backbone for more efficient Cre-dependent handle switching. So far, our screens using different Cre drivers showed strong correlation between independent screens, but no clear evidence of genes essential only in specific neuronal populations. This could be due to the genes included in the M1 library, which may not be optimally suited to detect population-specific vulnerabilities. It may also require longer screening durations to identify genes that impart differential susceptibility.

When applying CrAAVe-seq to new applications, factors such as sgRNA library size and viral tropism must be considered. Unlike CRISPR screens in cultured cell-based systems, where low MOI ensures that each cell mostly receives only one sgRNA, the complexity of CNS architecture and viral tropism in different brain regions makes controlling global MOI challenging, as each region will have a local MOI. In our studies, the MOI across the brain was generally 1 to 2, with a few areas exceeding 3. Despite this variability, the large screening library and large number of transduced neurons greatly buffers the potential for complex sgRNA interaction within the same cell, ensuring discernible phenotypes even when randomly paired with other sgRNAs. We found that reducing AAV library concentration significantly diminishes screen performance, indicating that transducing a large number of neurons is more critical than maintaining a low MOI. We also observed, through bootstrapping analyses, that a screen on smaller neuronal populations requires a larger number of mice. Future CNS screens must balance library size, target population(s) with consideration of dissected brain regions, and desired phenotypes, and we recommend using digital PCR and bootstrap analyses in optimizing for *in vivo* screens.

This platform can be immediately applied to existing neurodegeneration mouse models to systematically screen for modifiers of neuronal vulnerability in large brain regions like the cortex or focused populations, such as Purkinje neurons, where the abundance of internal granule cells might mask biological hits. Moreover, the persistent expression of AAV episomes in non-dividing cells^34^ makes CrAAVe-seq well-suited for aging studies. While our screen mainly identified gene perturbations reducing neuronal survival, future studies in neurodegenerative models could identify perturbations that rescue neuronal survival to reveal potential therapeutic targets. Beyond neurodegeneration, we conceive possible applications to study genetic factors that modify cell survival or growth/proliferation after traumatic brain injury and stroke.

Further, while our interest is in the brain, we believe this strategy will be useful for screening specific subpopulations of cells throughout the body. The growing toolkit of AAV capsids and Cre drivers allows for immediate application of this approach to study a wide array of different cell types. A minor limitation of using non-integrating AAVs is the gradual loss of AAV episomes during cell division, reducing the fraction of cells from which phenotypic data can be collected, particularly in proliferating tissues. Consequently, this approach would not be best suited to screen for phenotypes related to increased cellular proliferation. This is not a major issue for neurons, as they are postmitotic, and this still permits screening for genes that prevent neuronal death in disease models. For other proliferating cell types, integrating a transposase system into the host genome could help maintain sgRNA expression, as demonstrated previously^35^. While this would partly negate the advantage of recovering sgRNAs from episomes, the other advantages afforded by AAV (**Extended Data Fig. 1a**), including superior biodistribution and safety profile remain valuable.

CrAAVe-seq complements orthogonal *in vivo* CRISPR screening techniques but is especially well suited for probing specific cell populations with far larger sgRNA libraries for unbiased biological discovery. By taking advantage of widespread AAV transduction in the brain, CrAAVe-seq allows screening across millions of neurons per mouse, which was not feasible with prior options, and this can be further scaled by using multiple mice per screen. While CrAAVe-seq does not provide the same cell-type resolution as scRNA-seq-based Perturb-seq, CrAAVe-seq offers a practical and far less expensive approach for large-scale initial screens using broader libraries (>1,000s of sgRNAs vs. ∼10s of sgRNAs) and larger targeted cell populations (∼millions vs. 10,000s of cells). We are currently in developing a genome-wide library in the pAP215 backbone for future CrAAVe-seq applications. While our current work demonstrates the use of CrAAVe-seq to uncover modifiers of cell survival, the strategy can be expanded to other relevant phenotypes in the future. For example, delivering an AAV library *in utero* could be used to comprehensively profile the genes involved in the migration and differentiation of different cell types in development. As for previous screens in cultured cells, the use of reporters and fluorescent read-out in conjunction with flow cytometry will enable the identification of modifiers of a plethora of cellular phenotypes. Thus, CrAAVe-seq has the potential to accelerate the rate of biological and therapeutic discovery in relevant animal models while minimizing cost and animal use.

## Author contributions

BR, IVLR, NT, AP and MK contributed to the study’s overall conception, design, and interpretation, with input from the other authors. BR, IVLR and NT created the figures and BR, IVLR and MK wrote the manuscript, with NT and AP contributing further in Materials and Methods. RT, KM and JP contributed preliminary experiments using lentiviral injection of mice. BR, IVLR, AP, SDB, TS and LY conducted and analyzed all other experiments. NT performed bootstrapping analysis and wrote the CRISPR screen analysis pipeline.

## Acknowledgements

The authors would like to thank other members of the Kampmann Lab for discussions on data analysis (Olivia Teter, Steven Boggess, Emmy Li, and Reet Mishra). We also thank Abby Oehler and Rigo Roman-Albarran for help with imaging and discussing protocols. We thank Charlie Gersbach and lab for providing the LSL-dCas9-KRAB mouse line. For assistance and advice with sequencing, we would like to thank Lea Kiefer from the Daniele Canzio’s lab, Eric Chow and team at the UCSF Center for Advanced Technologies, and Angela Detweiler at the Chan Zuckerberg Biohub Network. For assistance and advice with imaging, we would like to thank Caroline Mrejen and team at the Innovation Core at the Weill Institute for Neurosciences at UCSF. We would also like to thank Alex Pollen, Mercedes Paredes, Daniele Canzio, Eric Huang, Dan Mordes, and Cathryn Cadwell and their labs for the use of numerous items of shared molecular biology and histology equipment.

This work was supported by the Ben Barres Early Career Acceleration Award from the Chan Zuckerberg Neurodegeneration Challenge Network (MK); a Tau Consortium Investigator Award (MK); NIH/NIA grant R01 AG082141 (MK); the UCSF-CIRM Scholars Training Program, CIRM grant EDUC4-12812 (IVLR); UCSF Hillblom/BARI Graduate Fellowship Award (IVLR); NIH/NINDS grant T32NS115706 (IVLR); NIH/NINDS grant R25NS070680 (BR) and 1K08NS133300 (BR); NIH/NIA grant RF1AG062234 (JJP); and NIH/NIA grant K99AG062776 (KM).

## Competing interests

BR, IVLR, AP and MK have filed a patent application on in vivo screening methods. MK is an inventor on US Patent 11,254,933 related to CRISPRi and CRISPRa screening, a co-scientific founder of Montara Therapeutics and serves on the Scientific Advisory Boards of Engine Biosciences, Alector, and Montara Therapeutics, and is an advisor to Modulo Bio and Recursion Therapeutics.

## Methods

### Animals

All mice were maintained according to the National Institutes of Health guidelines and all procedures used in this study were approved by the UCSF Institutional Animal Care and Use Committee. Mice were housed on a 12-h light/dark cycle at 22-25 °C, 50-60% humidity, and had food and water provided *ad libitum*. Mice were randomly assigned for the experimental groups at time of injection and both male and female mice were used. In accordance with approved protocol, mice were monitored post injection and if signs of distress appeared, mice were documented and euthanized promptly. The mice used in this study are LSL-dCas9-KRAB (LSL-CRISPRi) mice (B6;129S6-*Gt(ROSA)26Sor^tm2(CAG-cas9*/ZNF10*)Gers^*/J, RRID: IMSR_JAX:033066)^16^ and dCas9-KRAB mice (B6.Cg-*Igs2^tm1(CAG-mCherry,-cas9/ZNF10*)Mtm^*/J, RRID: IMSR_JAX:030000). A summary of the individual mice used for CRISPR screening and select *in vivo* experiments is provided in **Supplementary Table 1**.

### Plasmids

The screening vector pAP215 (**Fig. 1a**, fully annotated map in **Supplemental File 1**) was generated using the pAAV-U6-sgRNA-CMV-GFP plasmid as the starting backbone (Addgene plasmid # 85451, a gift from Hetian Lei)^36^. We replaced the sgRNA scaffold sequence with one from pMK1334 (Ref. ^1^) and the mU6 using a gene block (gBlock, IDT technologies) using a modified mU6 sequence as reported in Addgene plasmid #53187 (Ref. ^37^). The CMV-EGFP module was replaced with EF1a-2xmycNLS-tagBFP2 from pMK1334 by Gibson Assembly. The W3 terminator was cloned from Cbh_v5 AAV-saCBE C-terminal (Addgene plasmid # 137183, a gift from David Liu)^38^. The hGH was replaced by the SV40 from pMK1334. The Lox66 and Lox71 sequences and their orientation were copied from the pFrt-invCAG-Luc (Addgene plasmid # 63577, a gift from Ivo Huijbers)^39^ and were inserted along with the 175-bp intervening spacer as a gBlock.

Additional plasmids in this study include pENN.AAV.hSyn.HI.eGFP-Cre.WPRE.SV40 (Addgene # 105540, a gift from James M. Wilson), pENN.AAV.CamKII.HI.GFP-Cre.WPRE.SV40 (Addgene # 105551, a gift from James M. Wilson), CN1851-rAAV-hI56i-minBglobin-iCre-4X2C-WPRE3- BGHpA (Addgene # 164450, a gift from The Allen Institute for Brain Science & Jonathan Ting) (PMID: 33789083), and pAAV-FLEX-GFP (Addgene plasmid # 28304, a gift from Edward Boyden). The NLS-mScarlet and NLS-mNeonGreen AAV plasmids were generated by restriction cloning, replacing the GFP sequence in plasmid CAG-NLS-GFP (Addgene # 104061, a gift from Viviana Gradinaru)^17^, and replacing the NLS sequence with one from the pMK1334 plasmid.

### sgRNA cloning

We transferred the sgRNA sequences from our pooled mCRISPRi-v2 sgRNAs, subpools M1-top 5 (targeting Kinases, Phosphatases, and Drug Targets) and M3-top 5 (targeting the proteostasis network)^26^ into the pAP215 plasmid backbone. 20 µg of the library was digested with BstXI (Thermo Scientific, FD1024) and Bpu1102I (Thermo Scientific, FD0094). The guide-encoding inserts (84 bp) were resolved on a 4-20% Novex TBE gel (Invitrogen, EC62252BOX) and precipitated with GlycoBlue and sodium acetate. Inserts were washed with ethanol after precipitation and then eluted in DNase- and RNase-free water. 20 µg of the backbone vector, pAP215, was digested in parallel with BstXI and Bpu1102I, resolved on a 1% agarose gel, and purified from the gel (Zymo Research, D4001). The vectors and insert guides were annealed for 16 hrs overnight using T4 ligase (New England Biolabs, M0202L) at a 1:2 molar ratio of vector to insert, and then purified with sodium acetate and ethanol washing. After the final wash, a portion of the ligated library product was transformed into chemically competent *E. coli* (Takara, 636763) and 10 colonies were picked at random to ensure that each colony contained a unique sgRNA sequence. The remainder of the library product was electroporated into Mega-X competent cells (Invitrogen, C640003) and grown overnight, and a portion of the culture was plated to determine if a coverage of at least 250 colonies per guide was achieved, followed by growth of the remainder of the culture in 1 L of LB for 16 hrs and purification of the library using ZymoPURE II Plasmid Gigaprep Kit (Zymo Research, D4204).

The mouse chaperone targeting library was designed by selecting the mouse orthologs of a human chaperone targeting library that we previously developed^40^ and included 350 non-targeting control sgRNAs. Oligonucleotide pools were synthesized by Agilent, amplified by PCR, and cloned into the pAP215 backbone as above.

Individual sgRNAs were cloned in the pAP215 backbone digested with BstXI and Bpu1102I using annealed oligonucleotides with compatible overhangs. The protospacer sequences for the specific sgRNAs used in this study include sgCreb1 (GGCTGCGGCTCCTCAGTCGG), sgHspa5 (GAACACTGACCTGGACACTT’), sgRabggta (GCGGCGAACTCACCTGCTCA), and a non-targeting control (sgNTC) (GGATGCATAGATGAACGGATG).

### AAV packaging, purification, and titering

The pAP215-M1 library was packaged into AAV for transduction into neonatal mice as follows. Two 15-cm dishes were each seeded with 1.5×10^7^ HEK 293T cells (ATCC, CRL-3216) in 25 ml DMEM complete medium: DMEM (Gibco, 11965-092) supplemented with 10% FBS (VWR, 89510, lot: 184B19), 1% penicillin-streptomycin (Gibco, 15140122), and 1% GlutaMAX (Gibco, 35050061). The next day, 20 µg of pAdDeltaF6 (Addgene # 112867, a gift from James M. Wilson), 7 µg of pAP215-M1 library plasmid, 7 µg of pUCmini-iCAP-PHP.eB (Addgene # 103005, a gift from Viviana Gradinaru)^17^, and 75 µl of 1 mg/ml polyethenylamine (PEI) (Linear, MW 25,000, Polysciences, 23966) were diluted into 4 ml of Opti-MEM (Gibco, 31985062), gently mixed, and incubated at room temperature for 10 min. The PEI/DNA transfection complex was then pipetted drop-wise onto the HEK 293T cells. After 24 hours, the media was replaced with 27 ml of fresh Opti-MEM.

72 hours after transfection, AAV precipitation was performed as previously described^41^, with modifications. Cold 5× AAV precipitation solution (40% polyethylene glycol (Sigma-Aldrich, 89510) and 2.5 M NaCl) was prepared. The cells and media were triturated and collected (∼30 ml) into a 50 ml conical tube, followed by addition of 3 ml chloroform and vortexing for approximately 30 seconds. The homogenate was centrifuged at 3,000g for 5 min at room temperature, and the aqueous (top) phase was transferred to a new 50 ml conical tube and 5× AAV precipitation solution was added to a final 1× concentration, followed by incubation on ice for at least 1 hour. The solution was centrifuged at 3,000 × g for 30 min at 4°C. The supernatant was completely removed and the viral pellet was resuspended in 1 ml of 50 mM HEPES and 3 mM MgCl2, and incubated with 1 µl DNase I (New England Biolabs, M0303S) and 10 µl RNase A (Thermo Scientific, EN0531) at 37°C for 15 min. An equal volume of chloroform was added, followed by vortexing for 15 sec, and centrifuged at 16,000g for 5 min at RT; this step was repeated once. Using 400 µl at a time, the aqueous phase was passed through a 0.5-ml Amicon Ultra Centrifugal Filter with a 100 kDa cutoff (Millipore, UFC510024) by 3 min of centrifugation at 14,000g, followed by buffer exchange twice with 1× DPBS. Titering was performed by quantitative RT-PCR as previously described^42^ using primers listed in **Supplementary Table 4**.

To prepare AAV for testing in primary neuronal cultures (for longitudinal imaging and qRT-PCR), HEK293T cells were seeded into a 6-well format containing 1.5 ml of DMEM complete media. The cells were transfected with 1 µg pAdDeltaF6, 350 ng pUCmini-iCAP-PHP.eB, and 350 ng of AAV transgene using PEI as above. Approximately 48 hours after transfection, the cells and media were collected in 2 ml microfuge tube, 200 µl of chloroform was added to each tube, vortexed for 15 sec, and centrifuged at 16,000 × g for 5 min at room temperature. The aqueous (top) phase was transferred to a new tube and AAV precipitation solution was added to 1× dilution, and incubated on ice for at least one hour. The precipitated AAV was centrifuged at 16,000 × g for 15 min at 4°C, the supernatant was removed, the pellet was resuspended in 100 µl of 1× PBS, and centrifuged again at 16,000 × g for 1 min to remove excess debris, and the supernatant (purified virus) was transferred to a new microfuge tube. 10 µl purified virus was used per well in primary neuronal cultures in a 24-well format.

### Intracerebroventricular injection

Intracerebroventricular (ICV) injections were performed as previously described, with minor modifications^43^. Briefly, neonatal mice were placed on a gauze-covered frozen cold pack and monitored for complete cryoanesthesia. The scalp was gently cleaned with an alcohol swab. AAVs were diluted in 1× PBS with 0.1% trypan blue into a 2 µl final volume per mouse and loaded into 10 µl syringe (Hamilton, 1701). The syringe was equipped with a 33-gauge beveled needle (Hamilton, 7803-05, 0.5 inches in length). The needle was inserted through the skull 2/5 of distance of the lambda suture to the eye and to a depth of 3 mm to target the left lateral ventricle. Following a one-time unilateral injection, the neonate was placed on a warming pad for recovery and returned to the parent cage. The number of viral particles injected in each mouse is listed in **Supplementary Table 1**.

### sgRNA recovery and sequencing for CrAAVe-seq

Animals were euthanized using CO2, and their whole brains were removed and stored at -80°C. The sex of the mice was recorded prior to euthanasia.

#### Initial protocol for sgRNA recovery

This protocol for episome recovery was used in the following figures: Fig. 1d, M1 library screen in Fig 3, the hSyn1-Cre screen in Fig 5d, the hSyn1-Cre versus no Cre screens in Extended Data Fig. 6, and the screens in Extended Data Fig. 7. Each brain was placed in a PYREX 7 ml Dounce Homogenizer (Corning, 7722-7) with 2 ml of TRIzol (Invitrogen, 15596026) and thoroughly homogenized using the A pestle (0.0045 nominal clearance) for 10 or more strokes. 0.4 ml of chloroform was added, vigorously shaken for 30 seconds, and centrifuged at 12,000g for 15 min at 4°C. The aqueous phase (top) was collected and nucleic acids precipitated using 1 ml isopropanol, incubated on ice for 10 min, and centrifuged at 12,000g for 10 min at 4°C. The supernatant was discarded and the pellet was washed in 2 ml of 75% ethanol in DNase/RNase-free water and spun down at 7,500g for 5 min. The supernatant was then removed and the pellet was allowed to air dry for 10 mins, and then resuspended in 100 µl of DNase/RNase-free water and incubated with 1 µl of RNase A (Thermo Scientific, EN0531) at 37°C overnight. The sample was then column purified by Zymo DNA Clean & Concentrator-25 kit (Zymo Research, D4033) and eluted in 50 µl of DNase/RNase-free water to yield recovered viral DNA. The remaining RNAse-treated samples were considered recovered episomes for use in PCR below.

#### Optimized protocol for sgRNA recovery

An optimized protocol for episomal sgRNA recovery was used in the following figures: the M3 library screen in Fig. 3, the CaMKII-Cre screens in Fig 4., the hi56i-Cre screens in Fig. 5, and for the dPCR experiments in Fig 5. All steps in this protocol are the same as the above initial protocol except for two modifications. First, each brain was homogenized in 4 ml of TriZOL, phase separated using 0.4 ml chloroform, and the aqueous phase precipitated with 2 ml isopropanol, before resuspending in 100 µL of DNase/RNase-free water. Second, following overnight RNase A treatment as above, the sample was directly transferred to -20 ℃ without column purification.

#### PCR-amplification of sgRNAs and sequencing

The PCR was performed using Q5 High-Fidelity 2× Master Mix (NEB, M0492L). Each reaction contained 100 µL of recovered episomes, 110 µL of Q5 2× master mix, and 5.5 µL of each primer. For amplification of the AAV sgRNA libraries, the purified AAV was diluted 10-fold into H2O, and 1 µL of the diluted AAV was used as a template in a 100 µL PCR reaction. The reaction was distributed into PCR tubes at the maximum volume allowed by the PCR equipment.

The following PCR cycling conditions were used: 98°C 30s, (98°C 30s, 60°C 15s, 72°C 15s) × 23 cycles, 72°C 10min.

100 µL of each PCR reaction was purified using 1.1× SPRI beads (SPRIselect Beckman Coulter, B23317) and resuspended in 25 µL elution buffer (Machery Nagel, 740306). The purified products were pooled and sequenced on the Illumina HiSeq4000 at the UCSF Center for Advanced Technologies or the Illumina NextSeq2000. The amplification primers (with adapters) and custom sequencing primers are listed in **Supplementary Table 4**.

### Lentivirus packaging, purification, and injection

The pLG15 vector containing a non-targeting control sgRNA was packaged into lentivirus as previously performed^44^ by using PEI for transfecting 15 mg of the transfer plasmid and 15 mg of lentiviral packaging plasmids (containing 1:1:1 pRSV, pMDL, pVSV-G) into 1.0×10^7^ HEK293T cells cultured in a 15-cm dish in DMEM complete medium. 48 hours after transfection, the virus was precipitated from the media supernatant using Lentivirus Precipitation Solution (Alstem, VC100) and resuspended in 500 µl of PBS, and then further concentrated using the 0.5-ml Amicon Ultra Centrifugal 100 kDa Column. 1.8 µl of virus plus 0.2 µl of 1% Trypan Blue was injected by ICV for each neonatal mouse. Mouse brains were extracted on day 14 and sectioned coronally.

### Mouse cortical neuron primary cultures and immunocytochemistry

Neonates were briefly sanitized with 70% EtOH and decapitated using sharp scissors, and the brains were removed and placed into cold HBSS (Gibco, 14175095). The meninges were removed under a dissecting microscope, and the cortices were transferred to a 15-ml conical tube containing 10 ml of 0.25% Trypsin-EDTA (Gibco, 25200056) and incubated at 37°C for 30 min. The trypsin was removed and the brains were gently rinsed twice in 5 ml of DMEM complete media, followed by trituration of brains in 5 ml of DMEM complete media filtered through a 40 µm nylon cell strainer (Corning, 352340), and diluted into DMEM complete media in a volume as needed for plating. An equivalent of one brain was plated across each BioCoat Poly-D-Lysine 24- well TC-treated plate (Corning, 356414). The following day, day in vitro 1 (DIV1), the DMEM complete media was replaced with neuronal growth media composed of Neurobasal-A Medium (Gibco, 10888022), 1× B-27 Supplement minus vitamin A (Gibco, 12587010), GlutaMAX Supplement (Gibco, 35050079), and 1% penicillin-streptomycin (Gibco, 15140122). On DIV2, the cultures were further supplemented with cytarabine (AraC) to a final concentration of 200 µM (Thermo Scientific Chemicals, 449561000). The primary neuronal cultures were transduced with AAV on DIV4 and imaged starting 4 days after transduction.

### RNA isolation and quantitative RT-PCR

Using CRISPRi primary neurons at 11 days after transduction of AAV (sgNTC, sgHspa5, or sgRabggta), RNA was isolated with the Zymo Quick-RNA Microprep Kit (Zymo Research, R1050). Samples were prepared for qPCR in technical replicate in 10 µl reaction volumes using SensiFAST SYBR Lo-ROX 2× Master Mix (Bioline, BIO-94005), custom qPCR primers from Integrated DNA Technologies used at a final concentration of 0.2 µM and cDNA diluted at 1:20 by volume. qPCR was performed on a Bio-Rad CFX96 Real Time System C1000 Touch Thermocycler. The following cycles were run (1) 98°C for 3 min; (2) 95°C for 15 s (denaturation); (3) 60°C for 20 s (annealing/extension); (4) repeat steps 2 and 3 for a total of 39 cycles; (5) 95°C for 1 s; (6) ramp 1.92°C s^−1^ from 60°C to 95°C to establish melting curve. Expression fold changes were calculated using the ΔΔCt method, normalizing to housekeeping gene *Gapdh.* RT-qPCR primers are listed in **Supplementary Table 4**.

### Digital PCR

Digital PCR (dPCR) was performed to quantify percent of episomal sgRNAs with an inverted Handle sequence collected from mouse brains with or without hSyn1-Cre or hI65i-Cre. Episomal samples from the CrAAVe-seq post-RNase treatment step were diluted 1:50 in water prior to analysis. Two sets of primers were used: one for total sgRNAs (oIR020/oIR021, 132 bp amplicon) and one for sgRNAs with an inverted handle (oIR022/oIR023, 137 bp amplicon) with sequences in Supplemental Table 4. 15 µl dPCR reactions were prepared using 5 µl EvaGreen 3× PCR master mix (Qiagen, 250111), 1 µl of diluted episomal sample, 1.5 µl 10X primer mix (final primer concentration 0.4 µM each), and 7.5 µl nuclease-free water; 13 µl of this reaction was loaded into the microplate. dPCR was performed on a QIAcuity One 5-plex Digital PCR instrument (Qiagen) using 8.5k-partition, 24-well nanoplates (Qiagen, 250011). The thermal cycling conditions consisted of an initial 2 min at 95°C, followed by 40 cycles of 15 sec at 95°C, 15 sec at 60°C, and 15 sec at 72°C, with a final 5 min cooling step at 40°C. Image capture exposure was set to 250 ms. Samples included M1 library + hSyn1-Cre, M1 library only, and M1 library + hI56i-Cre, with four separate mice for each condition, analyzed for both total and inverted products. Analysis was performed on QIAcuity Software Suite (version 2.5.0.1). A common threshold of 75 RFU was set for all samples. Absolute concentration results for inverted sgRNAs were divided by total sgRNAs for each sample and multiplied by 100 to determine percent inverted sgRNAs.

### Mouse brain immunofluorescence staining

Whole brains were removed and fixed overnight at 4°C in 4% paraformaldehyde (Electron Microscopy Sciences, 15710) diluted in 1× PBS. The following day, the fixative was replaced with 30% sucrose dissolved in 1× PBS for at least 48 hours. Fixed brains were blotting dry, cut down the midline with a razorblade, and mounting into a cryomold (Epredia, 2219) using OCT compound (Sakura Finetek, 4583). To snap freeze, cryomolds were partially submerged in a pool of 2-propanol cooled by a bed of dry ice. Brains were sectioned in the sagittal plane at 40 µm on a cryostat (Leica, CM1950) with a 34° MX35 Premier+ blade (Epredia, 3052835). The resulting brain sections were stored free-floating in 1× PBS + 0.05% NaN3 at 4°C. When ready for staining, representative brain sections were wasted three times in 1× PBS and incubated in a 24 well plate at room temperature for one hour in blocking buffer: 10% goat serum (Gibco, 16210064), 1% BSA (Sigma-Aldrich, A7906), and 0.3% Triton X-100 (Sigma-Aldrich, T8787) diluted in 1× PBS. The brain sections were incubated in primary antibodies diluted in blocking buffer overnight at 4°C on a gentle shaker. The sections were washed three times in 1× PBS, then incubated in secondary antibodies for 2 hours at room temperature in the dark on a gentle shaker. Sections were washed three times in 1× PBS and moved to charged glass microscope slides (Fisher Scientific, 12- 55015). After PBS was removed, Fluoromount-G with DAPI mountant (Invitrogen, 00-4959-52) was added, and a No. 1.5 coverslip (Globe Scientific, 1415-15) was applied. Slides were dried at room temperature in the dark overnight and sealed with nail polish. For experiments without DAPI, ProLong Gold Antifade mountant (Invitrogen, P10144) was used instead. For experiments with Hoechst instead of DAPI, sections were lastly incubated for 15 mins in Hoechst 33342 (BD Pharmingen, 561908) diluted 2 µg/ml in 1× PBS, then washed 3 times in 1× PBS before mounting using ProLong Gold mountant.

The following primary antibodies were used: rabbit anti-CREB (1:1,000 dilution, clone: 48H2, Cell Signaling Technologies, 9197), rabbit anti-SOX9 (1:2,000 dilution, polyclonal, EMD Millipore, AB5535), guinea pig anti-NeuN (1:500 dilution, polyclonal, Synaptic Systems, 266004), alpaca FluoTag-Q anti-TagFP nanobody (reacts to mTagBFP2 but not eGFP, 1:500 dilution, clone: 1H7, Alexa647 pre-conjugated, NanoTag Biotechnologies, N0501-AF647-L. The following secondary antibodies were used: goat anti-rabbit IgG Alexa Fluor 488 (1:1,000 dilution, Invitrogen A-11034), goat anti-rabbit IgG Alexa Fluor 568 (1:1,000 dilution, Invitrogen, A-11011), goat anti-rabbit IgG Alexa Fluor 647 (1:1,000 dilution, Invitrogen, A-21245), goat anti-guinea pig IgG Alexa Fluor 488 (1:1,000 dilution, Invitrogen, A-11073), goat anti-guinea pig IgG Alexa Fluor 647 (1:1,000 dilution, Invitrogen, A-21450). All secondary antibodies were highly cross-absorbed.

### Microscopy, image segmentation, and analysis

Slides containing brain sections were imaged using a Zeiss AxioScan.Z1 with a Zeiss Colibri 7 unit, ×20/0.8 NA objective lens, 5-30 ms exposure, 1×1 binning and 25-100% intensity using 425- nm, 495-nm, 570-nm and 655-nm lasers, running ZEN version 2.6 software. The images were imported into QuPath (version 0.4.2) for analysis^45^. The raw CZI files are available on Dryad repository (see Data Availability).

To identify overlap between BFP, NeuN, and SOX9, a representative region of the cortex was outlined and the nuclei were segmented on the DAPI channel using the ‘Cell detection’ module without expansion of the nuclei to develop virtual cell boundaries. Classifiers were created to distinguish BFP^+^, NeuN^+^, and SOX9^+^ cells, and applied sequentially. Cells containing overlapping NeuN and SOX9 were considered to be neurons (as there was a low, but detectable signal in the SOX9 channel in all nuclei with this antibody) and only cells exclusively containing SOX9 were considered astrocytes. Similar segmentation on DAPI and sequential application of classifiers were used to examine overlap between nuclear BFP, mNeonGreen, and mScarlet signal in mice shown in Fig. 2d,e.

To evaluate CREB1 levels, a representative region was selected as indicated by specific brain regions, and the nuclei were segmented on the DAPI channel as above. The measurements for the segmented nuclei were exported. The mean fluorescence intensity for the anti-Creb1 channel was obtained selected by the top 200 nuclei of highest anti-mTagBFP2 fluorescence intensity. A representative region of brain stained with secondary antibodies only was selected to determine the background mean fluorescence intensity for that channel. The same segmentation was performed in mice injected with FLEX-GFP, with the top 2% and bottom 2% of GFP^+^ or BFP^+^ nuclei examined for CREB1 mean fluorescence intensity.

For mouse primary neurons transduced with AAV, live imaging was performed every other day using an ImageXpress Micro Confocal HT.ai High-Content Imaging System (Molecular Devices). The imaging chamber was warmed to 37°C and equilibrated with 5% CO2. The system used an Andor Zyla 4.5 camera with a Plan Apo ×10/0.45NA objective lens, an 89 North LDI laser illumination unit, 10-500 ms exposure time, 1×1 binning, and 10% laser intensity using 405-nm, 475-nm, and 555-nm lasers, running MetaXpress (version 6.7.1.157). Resulting images were imported into Cell Profiler (version 4.2.1)^46^ and analyzed using a custom pipeline. hSyn1-Cre- GFP+ nuclei were segmented using the ‘IdentifyPrimaryObjects’ module, with expected diameter 8-40 pixels, using an Adaptive threshold (size 50) and the Minimum Cross-Entropy method, with a 1.5 smoothing scale, 1.0 correction factor, and lower- and upper-bound threshold at 0.435 and 1, respectively. Segmented objects were exported, and counted in each field, then summed across all fields within a well to calculate the number of objects per well (n=29 fields per well, n=4 wells per condition), using a custom R script. This was repeated for each timepoint. Data was normalized to fluorescent intensity at day 8 (as before that day, fluorescence intensity increased linearly with time in all channels as cells manufactured fluorescent proteins) and percentage change was calculated for each well from day 8, for subsequent timepoints through day 16.

A similar protocol was used to analyze *Rabggta* knockdown data with some modifications. hSyn1- Cre-GFP^+^ nuclei were segmented using the ‘IdentifyPrimaryObjects’ module, with an expected diameter of 7-40 pixels, using an Adaptive threshold (size 50) and Minimum Cross-Entropy method, with a 1.3488 smoothing scale, 1.0 correction factor, and lower- and upper-bound threshold at 0.101 and 1, respectively. Segmented objects were exported and counted in each field, then summed across all fields within a well to calculate the number of objects per well (n = 29 fields per well, n = 3) using a custom R script. This was repeated for each timepoint. Data was normalized to fluorescent intensity at day 10 and percentage change was calculated for each well from day 10, for subsequent time points through day 26.

### CRISPR Screen Analysis

Computational analysis of screen data was carried out using a newly developed bioinformatics pipeline which is publicly available (see Code availability section). Raw sequencing results were mapped to the M1 protospacer library using ‘sgcount’. Briefly, ‘sgcount’ is a tool to match protospacers against a reference protospacer library with exact pattern matching. The resulting count matrices, containing guide and gene information along with count data for each sample, was used as input for subsequent analyses.

‘crispr_screen’ was used to perform differential sgRNA abundance analysis and gene score aggregation analysis. ‘crispr_screen’ is a reproduction of the original MAGeCK analysis but performs differential sgRNA analysis using a negative binomial as originally described in the study and not a truncated normal distribution as used in the current MAGeCK implementation. In brief, sgRNA abundances are median normalized across samples then a weighted linear regression (weighted ordinary least squares) is used to fit the log-variance to the log mean of the control samples (representing sgRNA abundances in the AAV library). The fit variance and mean are then used to parameterize negative binomial distributions for each sgRNA and a survival function or cumulative distribution function is used to calculate a p-value for sgRNA under- and over-abundance. We excluded any sgRNAs that were represented with fewer than 100 reads across the control AAV samples.

To calculate a gene-level aggregated metric across sgRNAs of the same gene group we established a novel algorithm, geopagg. We performed the following operations on the under- and over-abundance p-values in parallel. First, the differential abundance p-values for sgRNAs were corrected for multiple hypothesis correction using the Benjamini-Hochberg correction procedure to calculate a false discovery rate (FDR) for each sgRNA. Next, FDRs for sgRNAs belonging to the same gene grouping were collected and sorted ascendingly. We then calculated a weighted geometric mean FDR (*q_i_*) for each gene (*i* ∈ *N*) across the FDRs (*x_j_*) for sgRNAs within the gene group (*j* ∈ *M*).

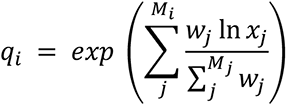

We calculated a weighted geometric mean to down-weight the relative impact of the first sgRNA within the group using a “Drop-First” weighting strategy. The first sgRNA (or top performing sgRNA) is down-weighted because we generally aim to capture genes with multiple high-performing sgRNAs. The weights for each gene grouping (*M_i_*) are defined as follows:

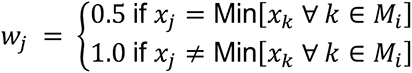

We also performed an aggregation of the log2-fold-changes in abundance (a gene’s phenotype score) of each sgRNA (*𝜑_j_*) within the gene group with an arithmetic mean:

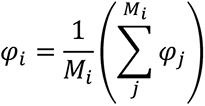

We then created random groupings of non-targeting control sgRNAs, which we denote as the *amalgam* gene set (*A*), to match the gene membership distribution of the input sgRNA library. This was performed by determining the membership size (number of sgRNAs) of each gene (*M_j_*) and sampling an equal amount of sgRNAs without replacement from the non-targeting controls. We next performed an identical calculation as above for each of the newly created *amalgam* genes.

We then calculated a ‘gene score’ for each gene and each *amalgam* gene within the dataset using the calculated weighted geometric mean of the FDR values *q_i_*) and their phenotype score (*𝜑_i_*).

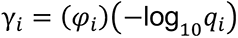

We next sort the gene scores (*γ_i_*) in an ascending order or a descending order for the under- and over-abundant tests respectively.

Finally, we calculate an empirical false discovery rate (*𝛿_i_*) by stepping through the weighted geometric mean (*q*) arrays and determining for each rank (*i*) how many *amalgam* genes (*g_i_* ∈ *A*) are preceding it. Because the true empirical false discovery rate will be zero for all genes preceding the first *amalgam* gene, we provide a non-zero score by constraining the reported false discovery rate to be the maximum of the empirical false discovery rate and the weighted geometric mean of that gene

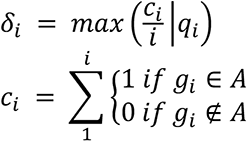

This empirical false discovery is further constrained for explicit monotonicity by requiring the current score to be greater than or equal to the previous one.

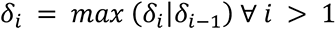

The geopagg algorithm is performed for the sgRNA under- and over-abundant p-values in parallel and the final scores for each gene are reported as the most significant of the two tests. 151 genes in the M1 library and 129 genes in the M3 library are targeted at two different transcriptional start sites by different sets of sgRNAs. These sets were evaluated independently with a label of P1 and P2 (e.g. GeneA_P1 and GeneA_P2). In cases where only one set is significant and labeled on a heatmap or volcano plot, the P1 or P2 label is not shown, but this information is included in **Supplementary Tables 2 and 3**.

### Bootstrapping Analysis

To estimate the required number of mice to effectively power a screen and assess the robustness of hit detection, we performed a bootstrapping analysis using a custom Python package (‘rescreener’). ‘crispr_screen’ was used to perform differential gene abundance analyses. The analysis was conducted as follows for each mouse experiment:

1. Full Dataset Analysis: The n=12 mice of the M1 library+CaMKII-Cre and n=11 mice of the M1library+hI56i-Cre were analyzed using the ‘crispr_screen’ tool to establish baseline results at an FDR of < 0.1, providing a list of hit genes from each full cohort
2. Bootstrapped Subset Analysis: From each cohort, multiple subsets of the treatment samples (mice) were randomly selected and analyzed:

a. Subset sizes ranged from 1 to the total number of treatment samples (mice)
b. For each subset size, 50 bootstrap replicates were generated by randomly sampling without replacement from the treatment samples (mice).
c. Each bootstrapped subset was analyzed using ‘crispr_screen’ with the same parameters as the full dataset analysis.
3. Overlap Assessment: For each bootstrap replicate, the overlap between its significant hits and those from the full cohort was calculated. Hits were defined as genes with an FDR of < 0.1.
4. Hit Recovery Analysis: For each gene identified as a hit in the full dataset analysis, we calculated the proportion of bootstrap replicates in which it was also identified as a hit. This proportion, termed the "recovery rate" was calculated as the number of times a gene was identified as a hit across all bootstraps divided by the total number of bootstrap replicates.

## Statistics and Reproducibility

No statistical methods were used to pre-determine sample sizes, but our sample sizes are similar to those reported in previous publications, as cited in the main text. Numbers of replicates are listed in each figure. No repeat measurements were made on the same samples. Data were assumed to be normally distributed except for instances within the ‘crispr_screen’ pipeline where geopagg uses a negative binomial distribution to calculate p-value. The ‘crispr_screen’ pipeline was used with FDR < 0.1 and controls for multiple comparisons using the Benjamini-Hochberg correction. For cell culture experiments, randomization was not performed because treatment groups of cells were derived from the same parent population of cells. Data collection and analysis were not performed blinded to the conditions of the experiments. No animals or data points were excluded from the relevant analyses. Major findings were validated using independent samples and orthogonal approaches.

## Data availability

All data are available from the corresponding author (MK), and will be made publicly available at the UCSF Dryad data repository upon publication (DOI: 10.5061/dryad.0k6djhb9t). There are no restrictions on data availability.

## Code availability

For CRISPR screen analysis, we developed a highly efficient analysis toolkit called ‘sgcount’ for sgRNA mapping and ‘crispr_screen’ for differential gene abundance testing. The sgRNA mapping utility (‘sgcount’, version 0.1.32) is also available on GitHub (https://github.com/noamteyssier/sgcount) and Zenodo (https://zenodo.org/doi/10.5281/zenodo.12774352). The differential gene abundance tool (‘crispr_screen’, version 0.2.8) is also available on GitHub (https://github.com/noamteyssier/crispr_screen/) and Zenodo (https://zenodo.org/doi/10.5281/zenodo.12774208) All bootstrapping analyses were performed using a custom python package (‘rescreener’, version 0.1.0) available on GitHub (https://github.com/noamteyssier/bootstrap_analysis_invivo_crispr_screen).

The R notebooks for analysis are available at https://kampmannlab.ucsf.edu/article/scripts-vivo-screening-manuscript.

The CellProfiler pipelines will be made available on request to the corresponding author (MK) and will also be submitted to the CellProfiler depository of published pipelines (https://cellprofiler.org/examples/published_pipelines.html) upon publication.

## Supplementary Materials

**Supplementary File 1: Annotated sequence of CrAAVe-seq plasmid pAP215.** A map of pAP215 is provided in a GenBank file format.

**Supplementary Table 1: Summary of mice injected with AAV across different studies** Lists all mice used in the study, time points, biological sex, and virus amounts. For mice used for CRISPR screens, the number of sequencing reads obtained from each sample is included.

**Supplementary Table 2: The CRISPR screen results of cohorts** Provides the results file of the ‘crispr_screen’ pipeline applied to the different cohorts of mice reported in this manuscript.

**Supplementary Table 3: CRISPR screen results for individual mice** Provides the knockdown phenotype (log2fc), pvalue, FDR, and Gene Score, obtained from the output of the ‘crispr_screen’ pipeline for all individual mice relative to input the AAV library, with each tab representing a different cohort of mice.

**Supplementary Table 4: List of primers used for sgRNA amplification, next generation sequencing, and RT-qPCR** Provides the primer sequences for sgRNA amplification, the custom sequencing primers for Illumina sequencing, and the primers used for RT-qPCR of Hspa5, Rabggta, and Gapdh.

**Supplemental Table 5: Values generated from bootstrap analyses for CaMKII-Cre and hi56i-Cre cohorts.** Three tabs each for bootstrap analyses of the CaMKII-Cre and hi56i-Cre cohorts reported in Fig. 6 are included. “Recovery by mouse number” tab shows the fraction of hits (frac_overlapping) that overlap with the hits of the full cohort when sampling different numbers of mice (subset). "All bootstraps” shows the total number (num_tests) and fraction (frac_tests) of all bootstraps that recover each hit of the full cohort. “Hits by subsample” shows the number (num_tests) and fraction (frac_tests) of bootstraps that recover each hit of the full cohort when sampling different numbers of mice (subset). Each test was run 50 times (replicate).

**Supplementary Video 1: Gross motor phenotypes in sgHspa5 + hSyn1-Cre injected mice** LSL-CRISPRi neonates were co-injected by ICV with AAV containing sgHspa5 (in pAP215) and AAV containing hSyn1-Cre. At 16 days post injection, mice displayed gross motor phenotypes, as recorded in the video. After video capture, mice were immediately euthanized as per IACUC protocol. Mice were littermates to those shown in Supplementary Video 2 and injected and recorded at the same timepoint.

**Supplementary Video 2: Lack of gross motor phenotypes in sgHspa5 only injected mice** LSL-CRISPRi neonates were co-injected by ICV with AAV containing sgHspa5 (in pAP215) and an equivalent volume of phosphate buffered saline. At 16 days post injection, mice displayed normal motor phenotypes, as recorded in the video. Mice were littermates to those shown in Supplementary Video 1 and injected and recorded at the same timepoint.

---

**Extended Data Fig. 1:**
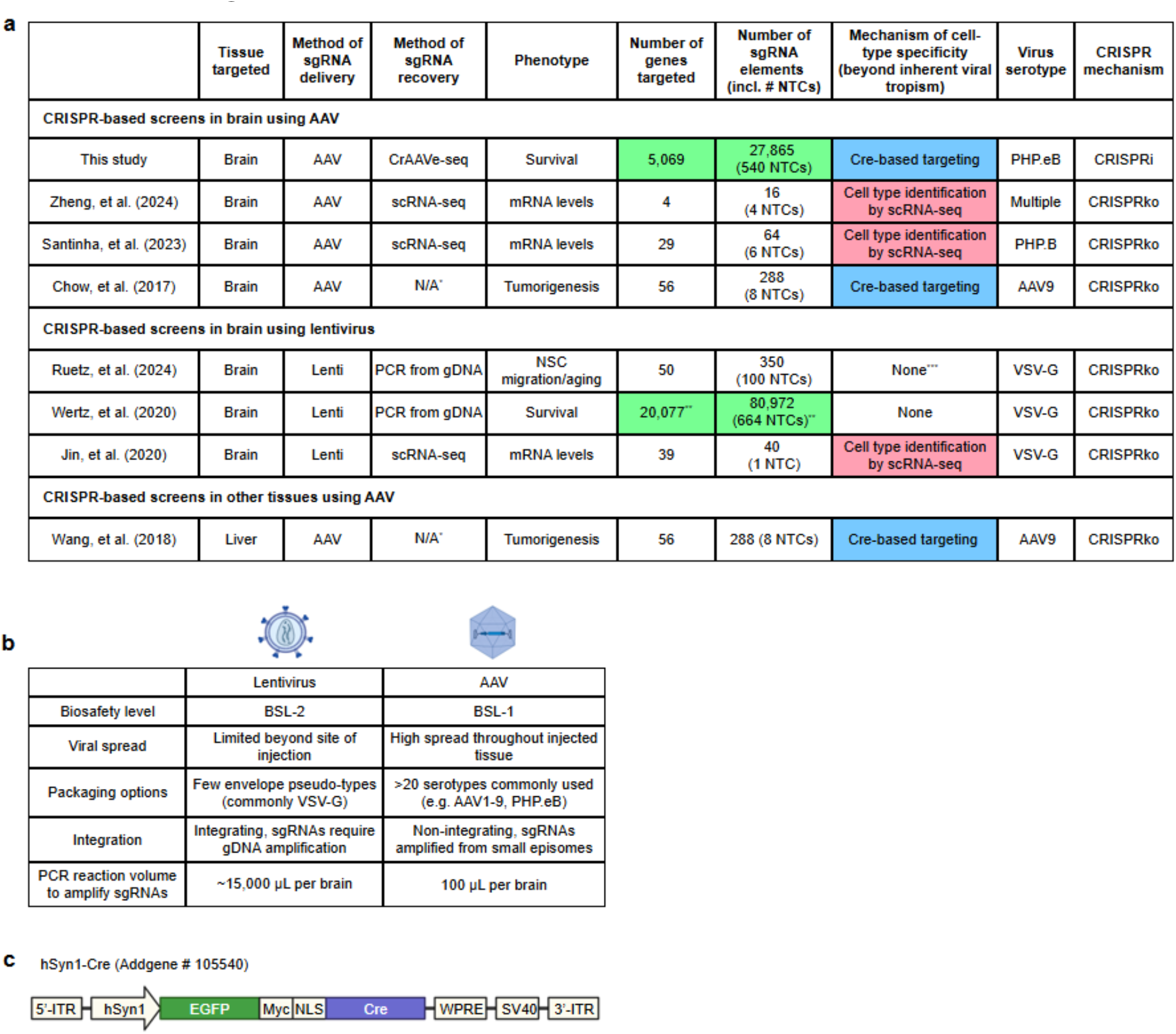
CrAAVe-seq enables simultaneously highly scalable and cell type-specific in vivo screens. a, Table summarizing published *in vivo* pooled screens of endogenous cells in the brain and AAV-based screen outside of the brain: Zheng et al.^9^, Santinha et al.^10^, Chow et al.^5^, Ruetz et al.^8^, Wertz et al.^6^, Jin et al.^7^, Wang et al.^47^. *Did not capture sgRNAs, instead performed mutagenesis screen; **Murine genome-wide Asiago library; ***A specific brain region containing migrated neural stem cell (NSCs) provided selectivity. CRISPRko: CRISPR knockout. Highlighted cells are large-scale sgRNA libraries (green), cell-type selectivity by Cre (blue), and single-cell RNA-sequencing (scRNA-seq, red). **b**, Table comparing different aspects of AAV and lentivirus. **c**, Simplified diagram of the hSyn1-Cre plasmid construct (Addgene # 105540).

**Extended Data Fig. 2.**
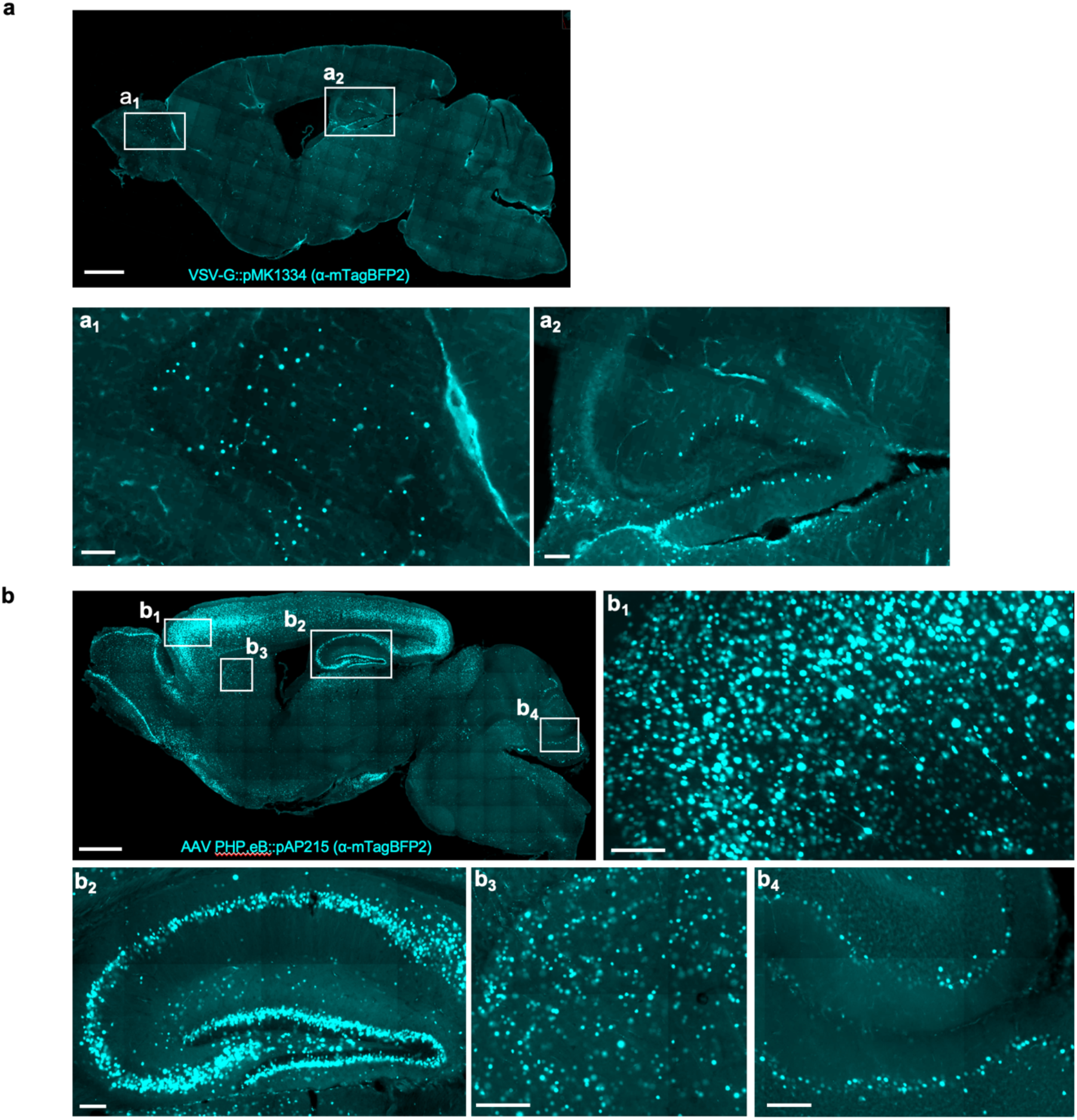
**a**, Sagittal section of mouse brain injected with sgRNA construct packaged in VSV-G-pseudotyped lentivirus (EF1a-NLS-mTagBFP2; pMK1334), stained with antibody recognizing mTagBFP2 at 3 weeks after ICV injection. Scalebar is 1 mm for low-magnification and 100 µm for subpanels (**a1**), (**a2**). **b**, Sagittal section of mouse brain injected with sgRNA construct packaged in AAV PHP.eB capsid (EF1a-NLS-mTagBFP2; pAP215), stained with antibody recognizing mTagBFP2 at 3 weeks after ICV injection. This condition is sgCreb1 + hSyn1-Cre (mouse 2), used in Fig. 2 and in Extended Data Fig. 4. Scalebar is 1 mm for low magnification and 100 µm for subpanels (**b1**), (**b2**), (**b3**), and (**b4**).

**Extended Data Fig. 3:**
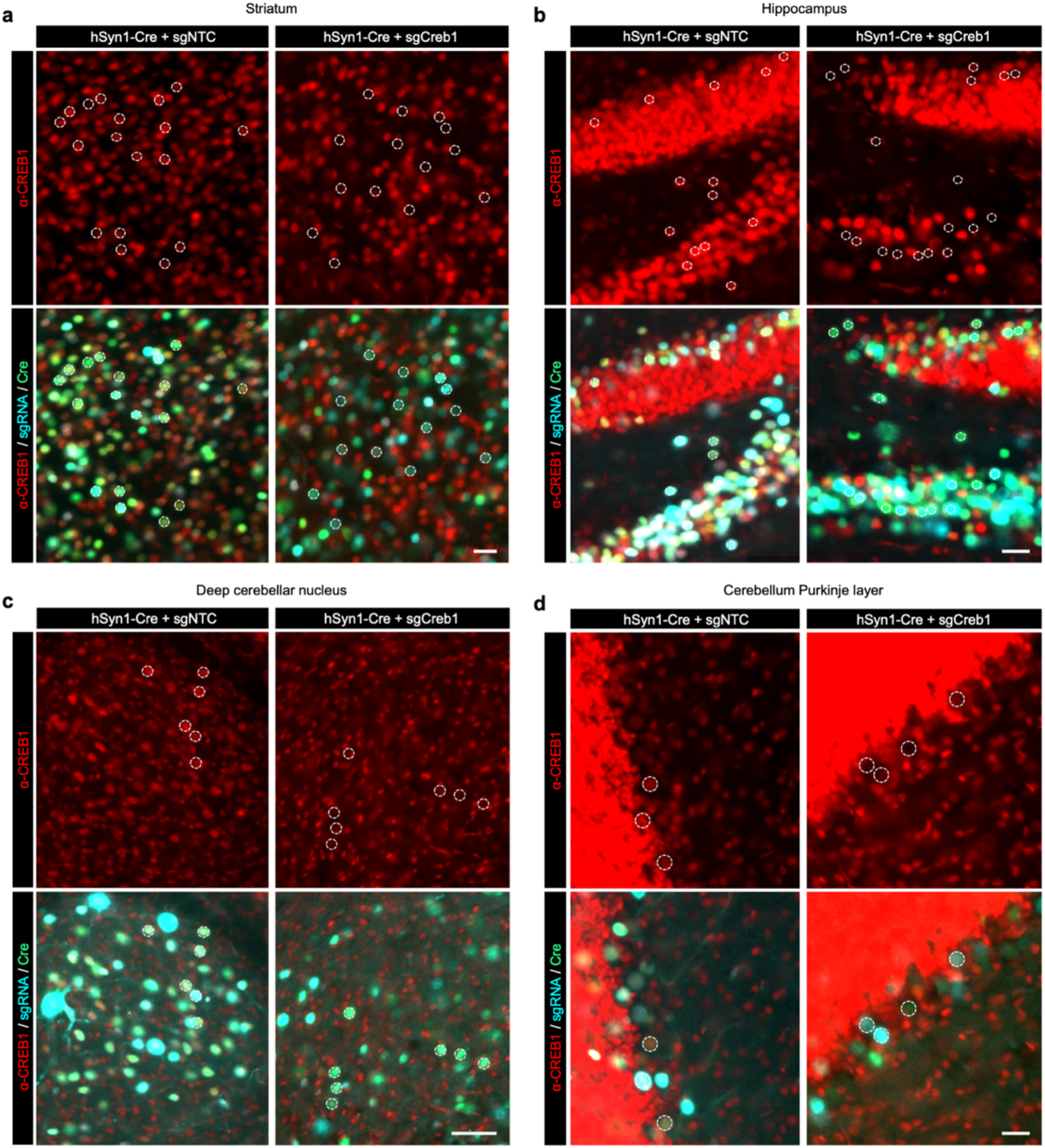
Cre-dependent CRISPRi knockdown of CREB1 in different brain regions of a mouse. Immunofluorescent staining for CREB1 (red) in an LSL-CRISPRi mouse 3 weeks after injection with PHP.eb::pAP215-sgNTC or -sgCreb1 (blue) along with PHP.eB::hSyn1-Cre (green), for different brain regions (**a**, Striatum, **b**, Hippocampus, **c**, Deep cerebellar nuclei, **d**, Cerebellum Purkinje layer). Representative nuclei containing both BFP and GFP are outlined with a dotted circle, with the same nuclei shown in the top and bottom panels of each brain region, highlighting markedly reduced CREB1 in double-positive nuclei with sgCreb1 as compared to sgNTC. DG: dentate gyrus; ML: molecular layer, Pkj: Purkinje cell layer (Pkj); IGL: internal granule layer. Data is quantified in Extended Data Fig. 4b.

**Extended Data Fig. 4:**
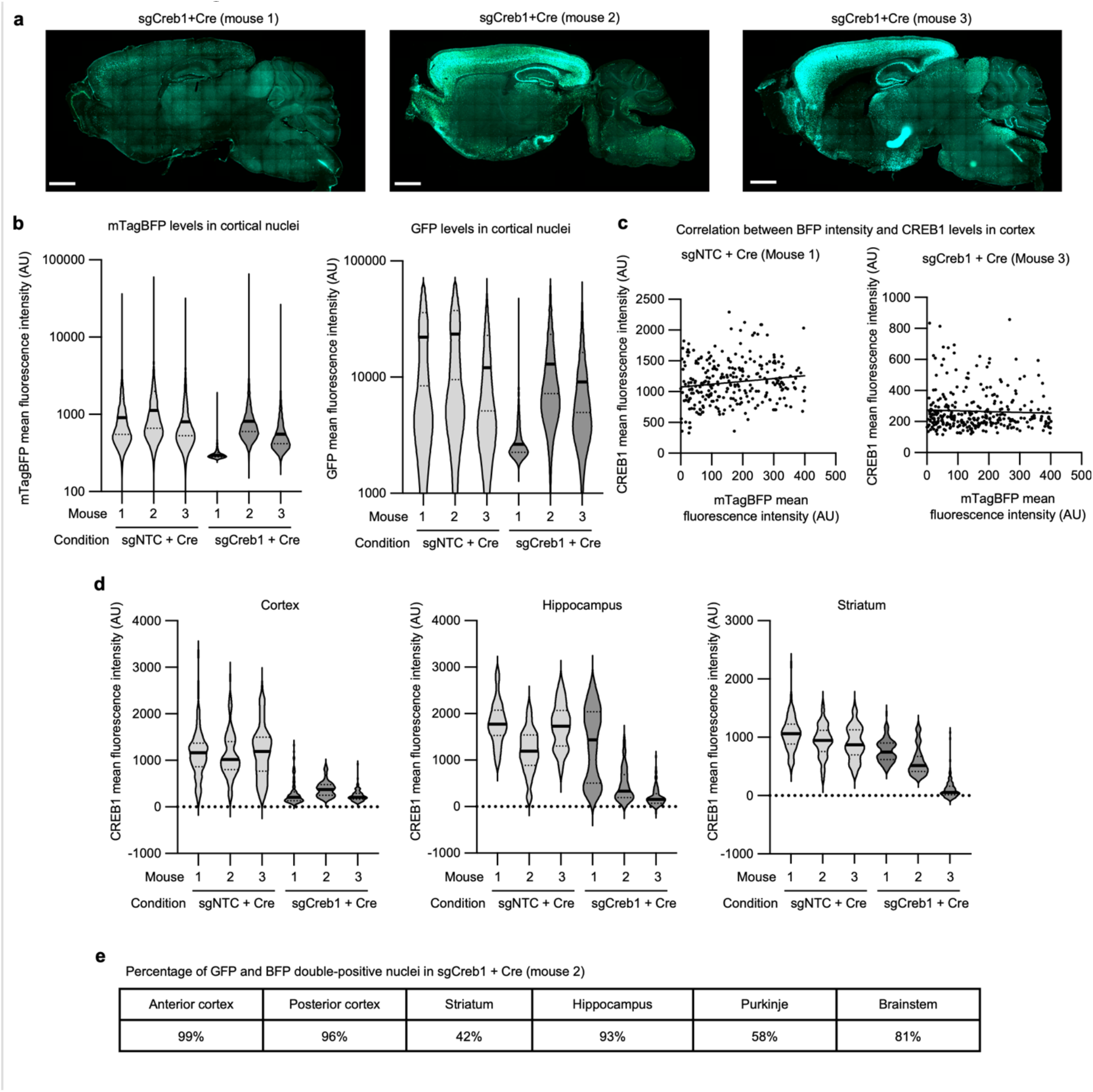
Quantification of CREB1 by brain region and by BFP intensity. **a**, Sagittal sections of the three mice injected with AAV pAP215 expressing sgCreb1 and AAV hSyn1-Cre (Cre) that are used in Fig. 2. Mouse 1 has a lower expression of both the sgRNA and Cre, potentially due to injection variability. **b**, Quantification of levels of BFP and GFP in a representative region of the cortex for sgNTC + Cre and sgCreb1 + Cre mice, n=3 each. **c**, CREB1 levels plotted against BFP levels in a representative region of cortex from two different mice, with trendlines generated by simple linear regression showing no correlation between BFP and CREB1 levels. **d**, Quantification of CREB1 levels in BFP^+^ nuclei of three different brain regions.

**Extended Data Fig. 5:**
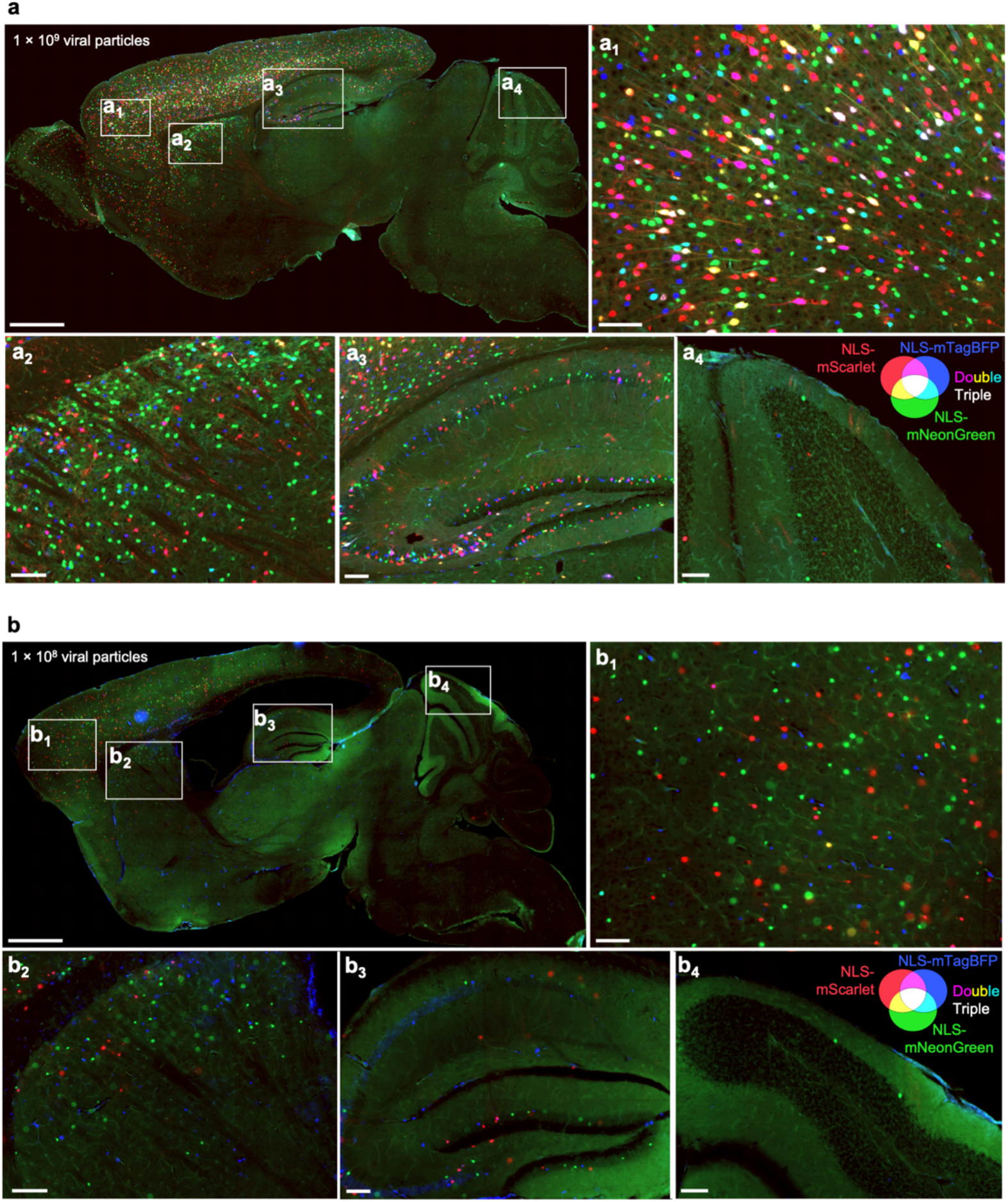
Distribution and extent of multiple infections with AAV injected at lower concentrations. Sagittal sections of mouse brains at 3 weeks after ICV co-injection of PHP.eB-packaged AAV expressing nuclear-localized mScarlet, mNeonGreen, or mTagBFP, at the indicated viral concentrations, which are (**a**) 10-fold or (**b**) 100-fold lower than show in Fig. 2. Scale bars: 1 mm for low-magnification views and 100 µm for subpanels.

**Extended Data Fig. 6:**
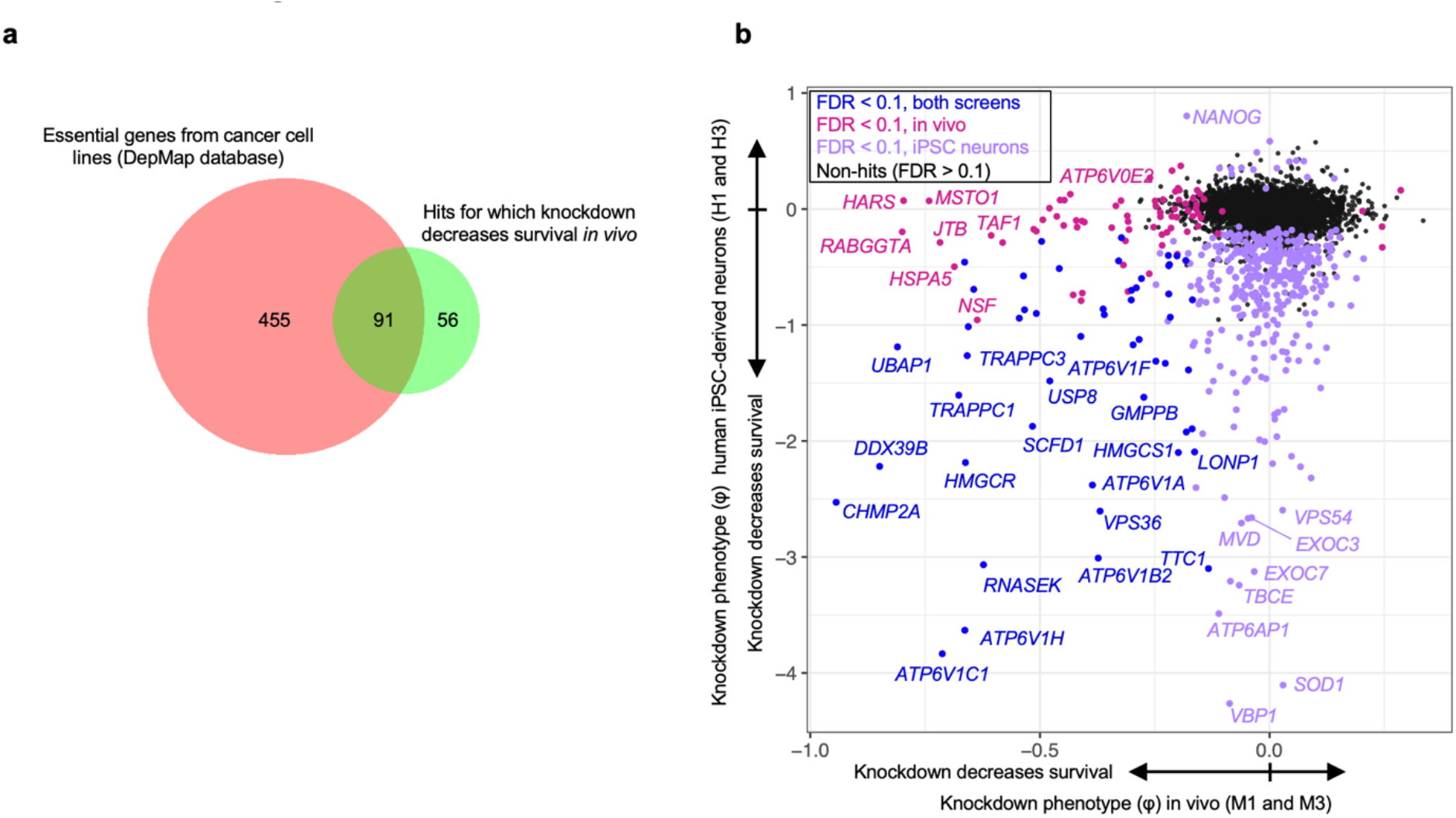
Overlap of hits of *in vivo* screens with essential genes of iPSC-derived neurons, and DepMap common essential genes. **a,** Venn diagram showing that 91 of the 147 hit genes for which knockdown decrease neuronal survival *in vivo* (Fig. 3) overlap with common essential genes of the injected libraries in cancer cell lines described in the DepMap database^27^, P<0.0001 by Fisher’s exact test, two-sided. **b,** Comparison of knockdown phenotype for genes between the *in vivo* neuronal survival screen of Fig. 3 and survival screens previously performed in iPSC-derived neurons by Tian et al.^2^.

**Extended Data Fig. 7:**
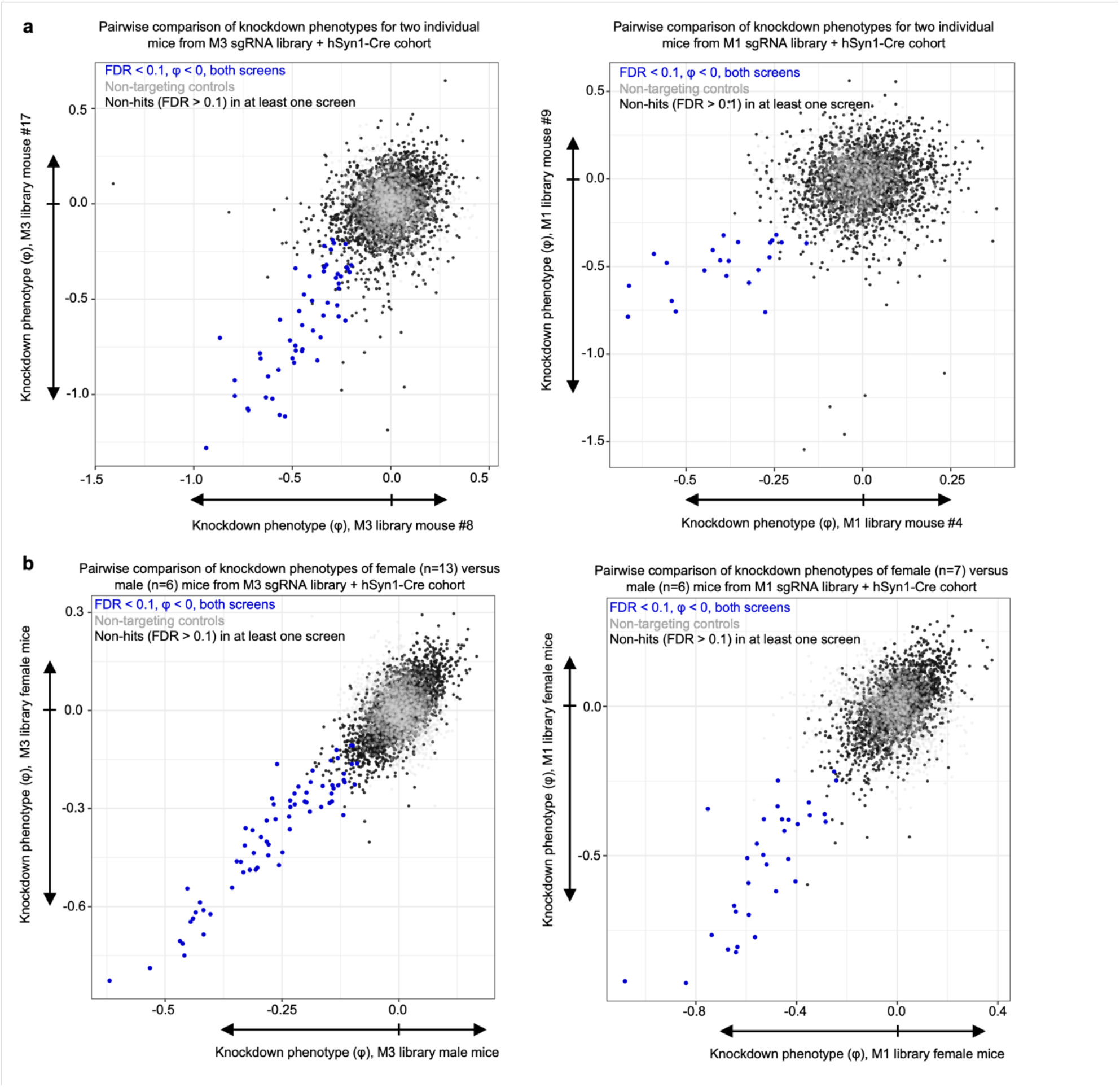
Pairwise comparisons of CRISPR screens by individual mice and by biological sex. **a**, Pairwise correlation of knockdown phenotypes between two individual mice injected with hSyn1-Cre and the M3 library (Left) and the M1 library (right). The two individual mice that called the greatest number of hits in each screen were selected for these comparisons. **b**, Pairwise correlation of combined knockdown phenotypes between male and female mice injected with hSyn1-Cre and the M3 library (left) and the M1 library (right).

**Extended Data Fig. 8:**
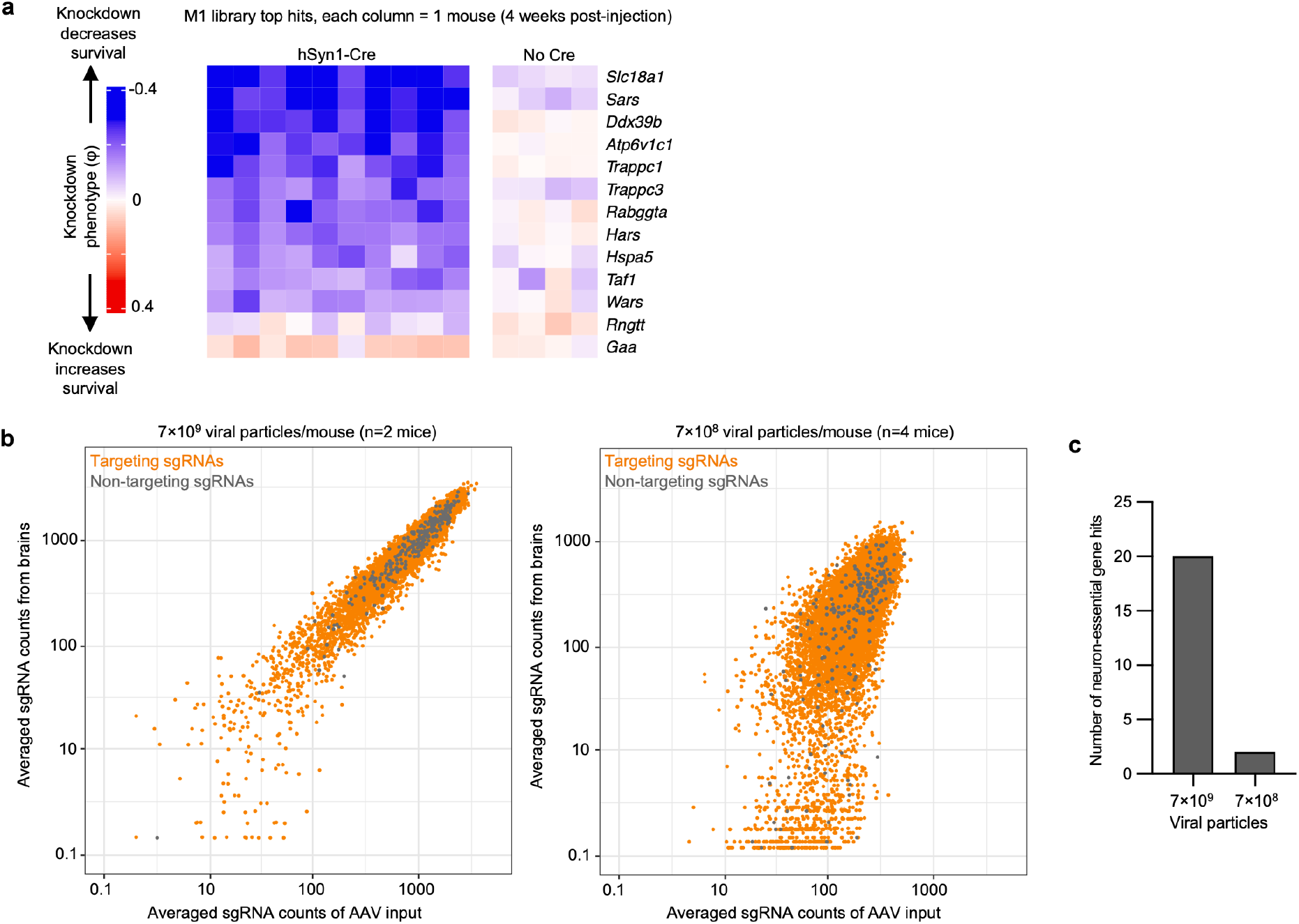
Cre expression and concentration dependency for screening performance. **a,** Heatmap of a subset of top hit genes in LSL-CRISPRi mice at 4 weeks after injection of PHP.eB::pAP215-M1 library, with or without co-delivery of PHP.eB::hSyn1-Cre. **b**, A cohort of LSL-CRISPRi mice were injected with either 7 x 10^9^ (n=2 mice) or 7 x 10^8^ (n=4 mice) viral particles of the PHP.eB::pAP215-M1 library in littermates. Normalized and averaged sgRNA counts from the brains are plotted against the sgRNA counts from the input AAV library, showing marked dropout of both targeting and non-targeting sgRNAs when using the lower concentration of virus. **c**, number of neuron-essential genes identified based on overlap with hits identified from the M1 library cohort in Fig. 3b in the mice with different concentrations of virus.

**Extended Data Fig. 9:**
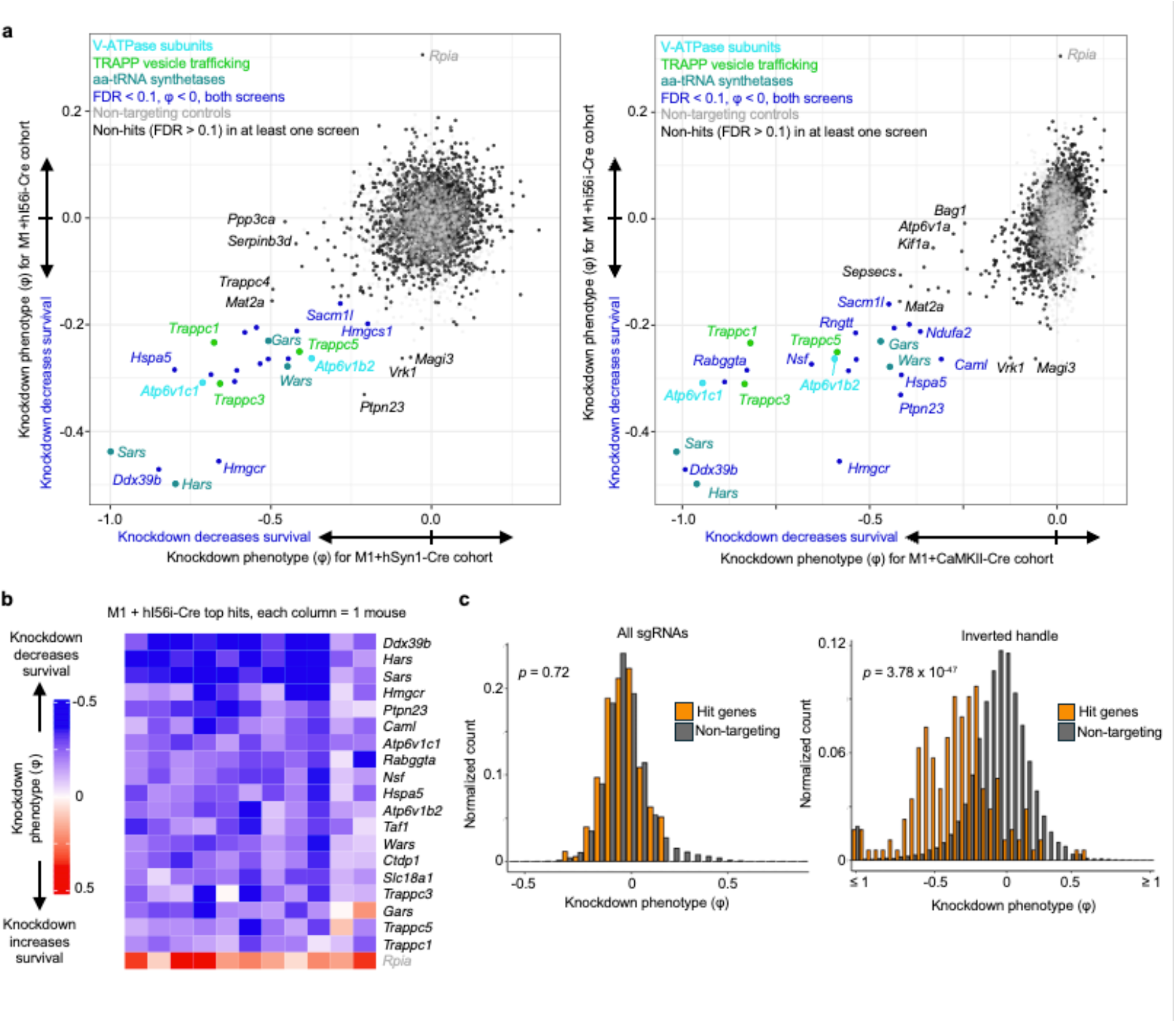
Knockdown phenotypes of essential genes identified by hI56i-Cre CRISPRi screen. **a,** Comparison of the knockdown phenotypes between the hI56i-Cre cohort and the hSyn1-Cre cohort of Fig. 3 (Left) and the CaMKII-Cre of Fig. 4 (Right). *Rpia* is labeled in gray to indicate that its effect is driven by only a single (but strong) sgRNA. **b,** Knockdown phenotypes plotted on a heatmap for top hit genes (rows, 19 with negative and 1 with positive knockdown phenotype) for all of the individual mice (columns). *Rpia* is shown in grey to reflect that its knockdown phenotype is driven by only one sgRNA. **c,** Histogram showing the distribution of gene knockdown phenotypes (φ) for either hit genes (orange), from (b), or for non-targeting control-based quasi-genes (grey) from the pAP215-M1 + hi56i-Cre screening cohort of mice shown in Fig. 5d. These plots compare the distributions for either all sgRNAs (left) or sgRNAs with the inverted handle (right). Knockdown phenotypes for each gene across the cohort were binned by 0.05 increments and normalized to the total number of genes (or quasi-genes) in that respective set. *P*-value calculated by asymptotic two-sample Kolmogorov-Smirnov test.

## Notes

### Summary of Updates

New computational analyses, systematic comparison with the previous literature, minor updates to figures.

